# CHOP: Haplotype-aware path indexing in population graphs

**DOI:** 10.1101/305268

**Authors:** Tom Mokveld, Jasper Linthorst, Zaid Al-Ars, Henne Holstege, Marcel Reinders

## Abstract

The practical use of graph-based reference genomes depends on the ability to align reads to them. Performing substring queries to paths through these graphs lies at the core of this task. The combination of increasing pattern length and encoded variations inevitably leads to a combinatorial explosion of the search space. We propose CHOP a method that uses haplotype information to prevent this from happening. We show that CHOP can be applied to large and complex datasets, by applying it on a graph-based representation of the human genome encoding all 80 million variants reported by the 1000 Genomes project.

## 1 Introduction

Pangenomes and their graphical representations have become widespread in the domain of sequencing analysis [1]. Part of this adoption is driven by the increased characterization of within species genomic diversity. For instance, recent versions of the human reference genome (GRCh37 and up), include sequences that represent highly polymorphic regions in the human population [2].

A pangenome can be constructed by integrating known variants in the linear reference genome. This way, a pangenome can incorporate sequence diversity in ways that a typical linear reference genome cannot. For example aligning reads to a linear reference genome can lead to an over-representation of the reference allele. This effect, known as reference allele bias, influences highly polymorphic regions and/or regions that are absent from the reference [3, 4]. By integrating variants into the alignment process, this bias can be reduced [5–7]. As a consequence, variant calling can be improved, with fewer erroneous variants induced by misalignments around indels, and fewer missed variants [8]. An intuitive representation for pangenomes are graph data structures, which are often referred to as population graphs [1, 9]. Population graphs can be understood as compressed representations of multiple genomes, with sequence generally represented on the nodes. These nodes are in turn connected by directed edges, such that the full sequence of any genome used to construct the graph can be determined by a specific path traversal through the graph. Alternatively, an arbitrary traversal through the graph will yield a mixture of genomes.

A key application for reference genomes is read alignment. Most of the linear reference read aligners follow a seed-and-extent paradigm, wherein exact matching substrings (seeds) between the read and a reference are used to constrain a local alignment. To efficiently search for exactly matching substrings (seeding), indexing data structures are used. The construction of these indexes generally relies on one of two methods: *k*-mer-based indexing, where all substrings of length *k* are stored in a hash-map along with their positions within the sequence; and sorting-based methods such as the Burrows-Wheeler Transform (BWT), where the reference sequence is transformed into a self-index that supports the lookup of exact-matching substrings of arbitrary length.

Existing indexing methods can be extended to population graphs, though this is challenging. Graphs can encode a variable number of genomes, which comes with an exponential growth of the number of paths in the graph as more variation is integrated. Arbitrary length sequence indexing is therefore not feasible, and indexing must be limited to shorter substrings to minimize combinatorial growth of the index. Additionally, sorting-based indexing methods that rely on suffix determination and ordering, are often unfeasible in graphs since there will be multiple valid node orderings.

Several approaches have been developed that perform read alignment onto population graphs using indexes that report all *k*-length paths in the graph. Early examples of this include: GenomeMapper [10], which builds a joint *k*-mer hash-map combining a collection of genomes to lookup seeds and subsequently align reads, using banded dynamic programming; BWBBLE [11], which linearizes the population graph using IUPAC encoding for SNPs and describes indels with flanking sequences as alternate contigs, after which it applies the BWT for indexing; In Satya et al. [12] they generate an enhanced reference genome from HapMap SNP-chip calls, wherein variants are encoded in read length segments used as alternative alignment targets alongside the reference genome. These methods are, however, orders of magnitude slower than linear reference genome aligners, or restricted to only small genomes. For instance, BWBBLE computes four times more suffix array intervals due to the expanded IUPAC alphabet. Moreover, these methods suffer from exponential growth in index space when variation density increases.

Increased scalability of population graph alignment has recently been demonstrated with Graphtyper [13] and the variation graph toolkit vg [14]. Graphtyper does so by first aligning reads to a linear reference sequence using BWA (as such there remains implicit reference bias), after which a graph-based alignment is performed on a much smaller set of unaligned or partially aligned reads. For this graph alignment it uses a *k*-mer hash-map of the population graph, wherein exponential growth in variation-dense regions is reduced through the removal of *k*-mers that overlap too many alternative sequences. The vg toolkit provides general solutions for working with population graphs. To efficiently query substrings, it utilizes GCSA2 indexing [15], an extension of the BWT for population graphs, which supports exact query lengths of up to 256 bp. Reads are aligned to graphs using a seed-and-extend strategy, returning subgraphs of the population graph to which reads are subsequently aligned using partial order alignment, a generalization of pairwise sequence alignment for directed acyclic graphs [16].

Since Graphtyper and vg index all possible paths in a population graph, they also cover complex regions where variation is dense. To deal with this, heuristics are utilized to prevent exponential growth. Either by removing *k*-mers that cross over more than a predefined number of edges, or by masking subgraphs shorter than a set number of bases. Techniques like these prevent exponential growth, but can completely remove complex regions in the graph, resulting in a loss of sensitivity in alignment. Furthermore, they contradict one of the main aims of population graphs, namely to address sequence variation in regions that are inaccessible through the application of a single reference sequence. An alternative solution that does not exclude complex regions, would be to constrain indexing by haplotype, so only *k*-mers observed in the linear genomes are encoded in the index. While the above heuristics are also used in vg, they recently also proposed the use of haplotyping. In vg such haplotyping is facilitated using the GBWT [17–19]. The GBWT is a graph extension of the positional Burrows-Wheeler transform [20], that can store the haplotypes of samples as paths in the graph, allowing for haplotype constrained read alignment. However, note that GCSA2 indexing will still require the evaluation of all paths in the graph, and will need graph pruning for complex graphs (any pruned paths can then be reintroduced into the graph with the GBWT).

We present CHOP, an alternative path indexer for graphs that incorporates haplotype-level information to constrain the resulting index. CHOP decomposes the graph into a set of linear sequences, similarly as in [12, 21], such that reads can be aligned by established aligners, such as BWA or Bowtie2 [22, 23], followed by the typical downstream analysis. We show that CHOP performs comparable to vg and more efficiently scales to human genomes with variation data of the 1000 Genomes Project [24, 25].

## 2 Results

Throughout, we consider population graphs constructed from variations called per sample (haplotype) with respect to a linear reference genome. These variations are encoded in the graph such that nodes represent sequences and edges represent observed consecutive sequences (Methods). CHOP facilitates read-to-graph mapping, which is presented in detail in the methods section. Briefly, CHOP transforms a population graph into a null graph (a graph devoid of edges) by a series of operations consisting of three steps: collapse, extend and duplicate, such that nodes in the null graph contain every substring of length *k* originating from the encoded original haplotypes in the population graph. Established aligners (here we have used BWA) can then be used to map reads to these null graph node sequences. Subsequently, these alignments can be projected back onto the population graph, given that the mapping of the node sequences in the null graph is known in the population graph (see Figure 1).

**Figure 1:**
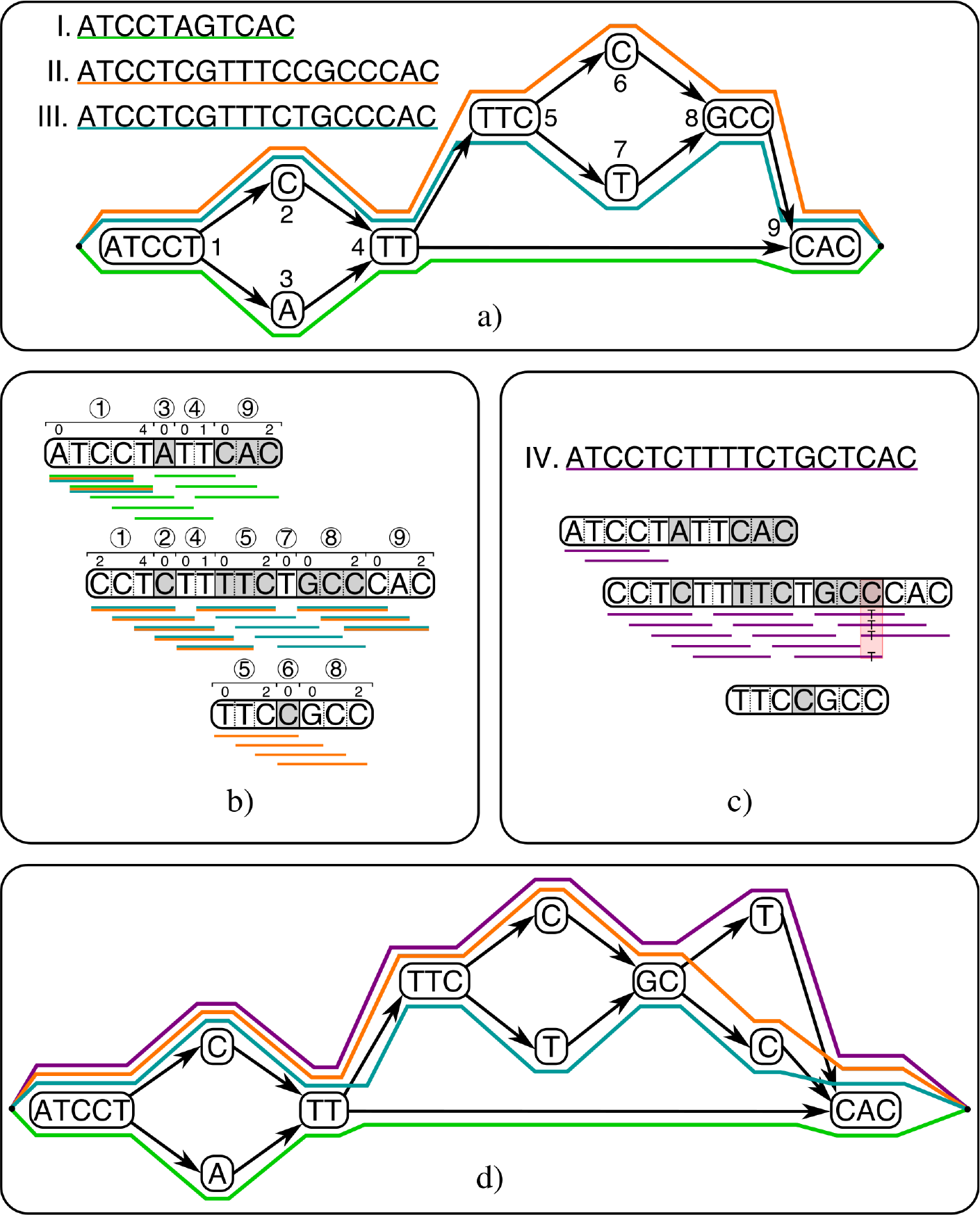
Schematic overview of how CHOP aligns reads to a population graph. a) As input, CHOP accepts a graphical representation of three distinct haplotypes (I, II, III). Colored paths through the graph identify underlying haplotypes. b) CHOP decomposes the graph into a null graph (a graph devoid of edges) for substrings of length 4 (Supplemental Figure S1 gives the full details about the decomposition). The obtained null graph contains three nodes, and the sequence that is defined on these nodes covers all substrings of length 4 that occur in the haplotypes encoded in the graph. Annotations above each node refer to intervals within nodes of the input graph. c) The reads (with length 4) from a new haplotype (IV) can be aligned to the null graph, consequently a mismatch can be called from the read pileup. d) Through the attached interval definitions that are assigned to the null graph, the novel variant can be positioned on node 8 of the original graph. Incorporating this variant results in a new graph.

### 2.1 Evaluation graph alignment

To evaluate CHOP, and its applicability in population graph alignment, we first performed tests on *Mycobacterium Tuberculosis* (MTB) using the read aligner BWA-0.7.15-MEM [22]. MTB represents a good model for population graphs, given the high accuracy of available assemblies, the tractable genome size (4.4 Mb), and the limited degree of variation. 401 variant call sets (VCF files) from different MTB strains (samples) were obtained from the KRITH1 and KRITH2 datasets [26, 27]. Variants were called with respect to the reference genome H37Rv, using Pilon-1.22 [28], and were filtered to exclude low quality variants. For graph construction we employed a leave-one-out strategy, wherein one sample was removed from the VCF file containing all 401 samples. The read set of the removed sample was subsequently used for graph alignment. This was repeated with 10 randomly selected samples. Corresponding single-end read sets were obtained from EBI-ENA (Supplemental Information 2). To investigate how introducing more variation influences graph alignment, we progressively incorporated more samples (from the complete set) into the constructed graphs. With up to 17,500 variants in the 400 sample graph (rate of variation growth is shown in Supplementary Figure S2).

As the ground truth of genomic positions in the read set data are unknown, we evaluated alignments based on the following criteria: number of mismatches, insertions, deletions, clipped bases, unaligned reads, and perfectly aligned reads (definitions in Supplemental Information 4). These criteria allowed us to inspect the behavior of different read aligners. In order to avoid bias induced by multiple possible alignments for a single read, we only considered primary alignments.

To evaluate our haplotype-based approach, we compared CHOP to vg-1.12.1 with haplotyping (denoted as vg+GBWT) and without. The vg toolkit provides general solutions for population graphs which include graph construction, indexing, and read alignments. CHOP was set to report 101-length haplotype paths (equivalent to the read length), and used default parameters with BWA-MEM. Vg was set to index all 104-length paths (*k* = 13, 3 doubling steps), to most closely reflect the settings of CHOP.

Because CHOP uses BWA as an aligner while vg has its own internal aligner, differences based on the aligner and not the indexing algorithm may occur. To understand aligner and parameter induced differences, we first summarized the results of the 10 hold-out samples on the linear reference genome, shown in Table 1 for BWA and vg. Both aligners resulted in nearly the same number of perfectly aligned reads. However, alignments with vg resulted in fewer unaligned reads (−22.30%) and more mismatches (+4.01%) than BWA. We attribute this difference to an increase in sensitivity by which vg aligns reads. This is reflected by the increase in clipped bases (+22.79%), inserted bases (+29.36%), and deleted bases (+34.53%), which allows vg to map shorter read fragments.

**Table 1:** Mean of alignment results across all 10 hold-out sample alignments to 1) the reference genome H37Rv (H37Rv columns) and 2) the 400 TB genomes graph (Graph columns) for both CHOP/BWA and vg with and without haplotyping to align the reads (note that when aligning only to H37Rv, CHOP is not used).

| Alignment criteria | All TB hold-out samples - Read length = 101 |  |  |  |  |
| --- | --- | --- | --- | --- | --- |
|  | BWA | CHOP/BWA | vg | vg | vg + GBWT |
|  | H37RV | Graph (n=400) | H37RV | Graph (n=400) | Graph (n=400) |
| Reads aligned | 6,160,920 | 6,162,033 (+0.018%) | 6,241,270 | 6,245,907 (+0.074%) | 6,244,004 (+0.044%) |
| Reads unaligned | 360,236 | 359,123 (-0.309%) | 279,886 | 275,249 (-1.657%) | 277,152 (-0.977%) |
| Reads perfectly aligned | 4,048,774 | 4,142,052 (+2.304%) | 4,048,774 | 4,153,217 (+2.580%) | 4,153,124 (+2.577%) |
| Bases aligned | 596,380,132 | 596,611,260 (+0.039%) | 599,244,753 | 599,601,399 (+0.060%) | 599,528,267 (+0.047%) |
| Bases unaligned | 62,191,423 | 61,960,355 (-0.372%) | 59,307,655 | 58,949,429 (-0.604%) | 59,023,102 (-0.480%) |
| Bases unaligned from clipped reads | 22,349,569 | 22,380,472 (+0.138%) | 27,442,533 | 27,690,552 (+0.904%) | 27,575,464 (+0.484%) |
| Bases mismatched | 3,458,029 | 3,308,480 (-4.325%) | 3,596,667 | 3,458,707 (-3.836%) | 3,455,296 (-3.931%) |
| Bases inserted | 65,210 | 65,151 (-0.090%) | 84,358 | 85,938 (+1.874%) | 85,397 (+1.232%) |
| Bases deleted | 52,272 | 51,165 (-2.118%) | 70,324 | 72,082 (+2.500%) | 70,347 (+0.033%) |
| Non-primary alignments | 246,092 | 246,540 (+0.182%) | 539,309 | 724,904 (+34.414%) | 724,613 (+34.360%) |
| Time (seconds) | 533 | 721 | 10,711 | 4,457 | 4,540 |

Using these measurements as a baseline, read to graph alignments were compared between CHOP/BWA and vg. The different graph constructions of CHOP and vg were found to have minimal effect on alignments as shown in Supplemental Figures S3 and S4. Figure 2 shows the increase in perfectly aligned reads using both CHOP/BWA and vg as more samples are incorporated into the graph (similar plots for the number of unaligned reads and mismatches can be found in Supplemental Figures S5 and S6). Table 1 shows the alignment results for the TB graph with 400 samples.

**Figure 2:**
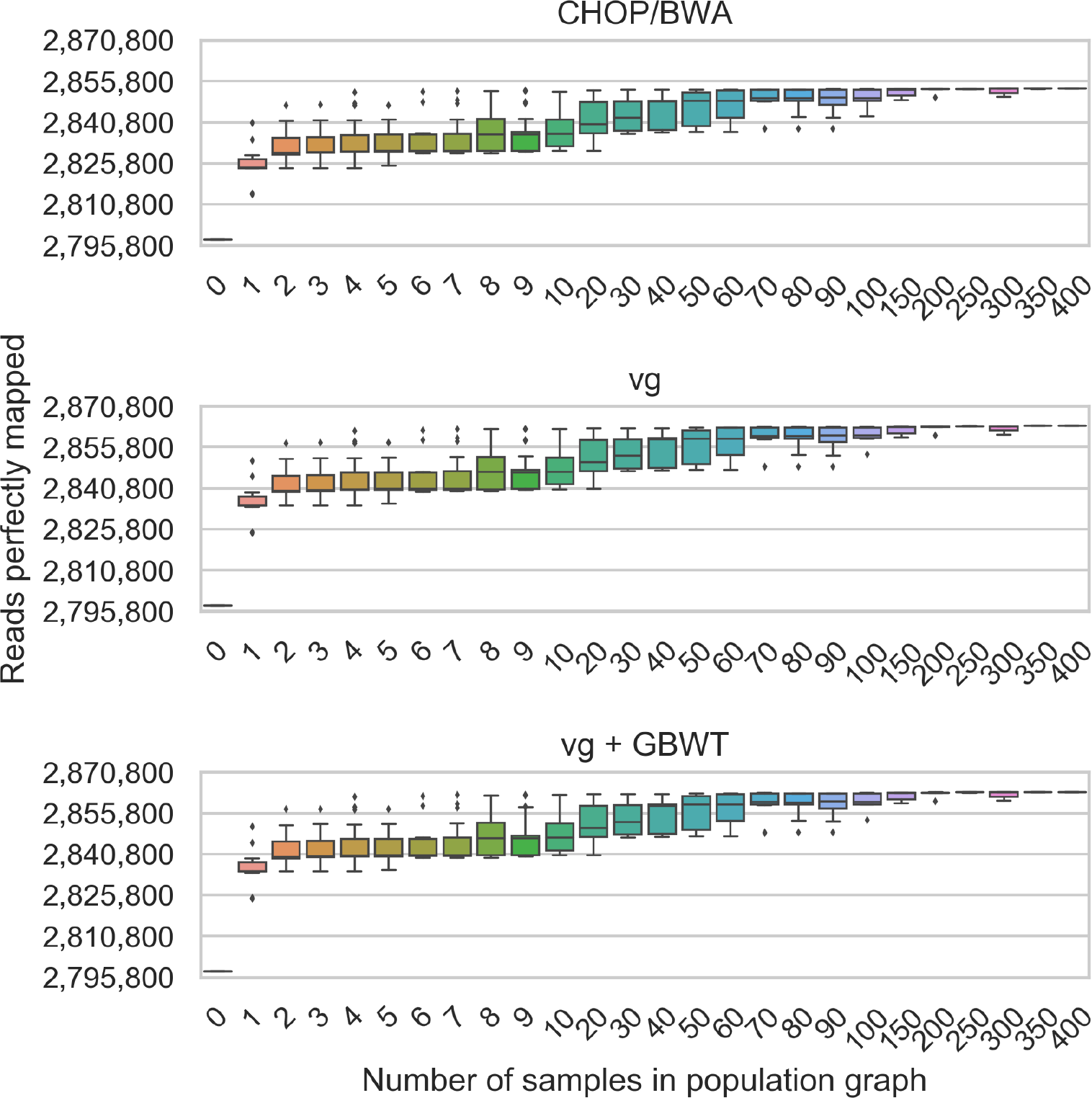
Perfectly aligned read count for SRR833154 alignments to different sized population graphs, containing between 0 (only H37Rv, the linear reference) and 400 samples for both, when using CHOP/BWA and vg with and without haplotyping to align reads to the graph.

Figure 2 shows that incorporating more variation from samples into the population graphs increases the number of aligned bases, which is further demonstrated in Supplemental Figures S5 and S6. Spread is a consequence of sampling when building the population graphs, where samples that are closely related to the hold-out sample will yield greater improvement than distantly related samples, further demonstrated by the reduction of spread as sampling size increases.

By comparing vg and vg+GBWT the effects of haplotyping can be observed, noting a drop in the number of aligned reads. This is to be expected as the index space has been constrained to only the haplotypes.

The baseline alignments to H37Rv already highlighted that the aligners perform differently. However, throughout the course of the experiments, nearly all alignment criteria show the same trend for both CHOP/BWA and vg. The exception being the number of unaligned reads, which steadily decreases with vg, while this is not as pronounced using CHOP/BWA. This difference can partially be attributed to the use of unconstrained path indexing compared to haplotyped indexing (difference is smaller with vg+GBWT), the remainder being caused by differences in aligner sensitivity.

### 2.2 CHOP scales to *Homo sapiens*

To further evaluate scalability and sensitivity of CHOP, we used chromosome 6 (170 Mb) of the GRC37 assembly in combination with the 1000 Genomes Phase 3 variation dataset [24, 25]. The constructed graph has 14,744,119 nodes and 19,770,411 edges and encodes a total of 5,023,970 variants (4,800,102 SNPs, 97,923 insertions, and 125,945 deletions). Note that the variation set included diploid phasing of 2,504 individuals, which was incorporated into the graph as 5,008 paths (2 paths per sample), and additionally one path that represents the reference genome. Within the population most variation (58.42%) is shared between at least two or more samples (Supplemental Figure S7). We used 15 single-end read sets from the 1000 Genomes project for the graph alignments (Supplemental Table S2), that were filtered to include only reads aligned to chromosome 6 or that could not be aligned anywhere on the genome (average read set size of 3,026,069).

CHOP was set to report 100-length paths through the graph to match the read length, which yielded 11,359,686 nodes in *G^E^*. The memory usage and time taken for indexing was dominated by CHOP, BWA indexing accounting for only 6.95% of indexing time and a fraction of memory required. We attempted indexing with vg and vg+GBWT for paths up to 104 bp (*k* = 13, 3 doubling steps), but this did not finish due to memory constraints (500 GB). Instead, doubling was lowered to 2, and paths up to 52 bp were indexed. By incorporating haplotyping in vg, the indexing requires substantially more time (6x longer), than indexing without haplotyping, whereas memory usage remains constant. The read sets were aligned to both the linear reference of chromosome 6 and the graph representation, using either CHOP/BWA, vg, and vg+GBWT, which is summarized in Table 2.

**Table 2:** Mean of alignment results from 15 samples from the 1000 Genomes data when aligning to 1) the reference genome sequence of chromosome 6 (column GRC37), and 2) the population graph created from the 5,008 haplotypes, for both CHOP/BWA and vg with and without haplotyping.

| Alignment criteria | 1000 Genomes samples - Read length = 100 |  |  |  |  |
| --- | --- | --- | --- | --- | --- |
|  | BWA | CHOP/BWA | vg | vg | vg + GBWT |
|  | GRC37 | Graph (n=2504) | GRC37 | Graph (n=2504) | Graph (n=2504) |
| Reads aligned | 2,542,399 | 2,543,522 (+0.044%) | 2,684,925 | 2,726,051 (+1.532%) | 2,717,972 (+1.231%) |
| Reads unaligned | 483,670 | **482,548 (-0.232%)** | 341,144 | **300,018 (-12.056%)** | **308,098 (-9.687%)** |
| Reads perfectly aligned | 1,794,564 | 1,977,952 (+10.219%) | 1,807,158 | 1,993,967 (+10.337%) | 1,993,469 (+10.310%) |
| Bases aligned | 251,122,992 | 251,516,725 (+0.157%) | 254,518,323 | 255,911,471 (+0.547%) | 255,578,370 (+0.416%) |
| Bases unaligned | 51,439,949 | 51,070,534 (-0.718%) | 48,029,518 | 46,654,663 (-2.863%) | 46,995,159 (-2.154%) |
| Bases unaligned from clipped reads | 1,801,947 | **1,846,687 (+2.483%)** | 12,716,162 | **15,699,089 (+23.458%)** | **15,245,035 (+19.887%)** |
| Bases mismatched | 1,270,981 | 969,087 (-23.753%) | 1,198,917 | 953,800 (-20.445%) | 940,371 (-21.565%) |
| Bases inserted | 43,979 | 19,661 (-55.296%) | 59,078 | 40,786 (-30.962%) | 33,391 (-43.480%) |
| Bases deleted | 61,659 | 32,355 (-47.526%) | 73,131 | 44,555 (-39.075%) | 41,040 (-43.882%) |
| Time alignment (seconds) | 544 | 1,807 | 19,996 | 10,436 | 10,871 |
| Memory alignment (MB) | 412 | 5,534 | 837 | 3,296 | 4,389 |
| Time indexing (seconds) | 186 | CHOP: 43,625<br>BWA: 3,256 | 37 | 5751 | 33,619 |
| Memory indexing (MB) | 245 | CHOP: 56,969<br>BWA: 3,813 | 269 | 45,670 | 45,868 |

We observed the same improvement of moving to a graph representation as in MTB, although more extensive, given that more variants, including indels, are incorporated into the graph.

Given the different path lengths used, time cannot directly be compared between CHOP/BWA and vg. Nevertheless, it is unclear why vg took substantially more time to align than CHOP/BWA, especially when mapping to the flat reference. Differences (relative to MTB) between vg and vg+GBWT have become more prominent, given that more samples are incorporated within the graph.

We observed substantial differences between CHOP/BWA, vg, and vg+GBWT with respect to the decrease of unaligned reads −0.232% versus −12.056% and −9.687%, and the increase in read clipping +2.463% versus 23.458% and 19.887%, respectively. To evaluate this aligner induced difference, we extracted all reads that aligned exclusively onto the graph, which amounted to 21,661 reads in CHOP/BWA and 616,900 in vg. Supplemental Figure S8 displays the distribution of the number of aligned bases for each of those reads. Nearly all (97.61%) of the newly aligned reads by vg had a length of between 15 to 30 bases, either induced by clipping or extensive base insertion/deletion. However, from 30 bases and up, the aligners display very similar profiles, with a comparable number of newly aligned reads. At 69 bp both aligners display a peak, the newly mapped reads corresponding to this peak all map to the same region in the graph. This region closely resembles human mitochondrial DNA, which was excluded from the initial reference alignments. This has led to an increased number of unmapped mitochondrial sequencing reads in the dataset that were mapped to the graph (Supplemental Information 9 for more details).

Finally, we performed alignments to the full human genome. We constructed graphs of each chromosome encoded with the variants as reported by the 1000 Genomes project. Cumulatively these graphs have 248,677,280 nodes, 33,3561,973 edges and encode a total of 84,745,123 variants (81,382,582 SNPs and 3,362,541 indels). We indexed the graphs with both CHOP for 100-length paths and with vg+GBWT for 52-length paths, the peak memory usage and time required for indexing is reported in Figure 3. Note that chromosomes 1, 2, 11, and X could not be indexed with vg+GBWT, due to graph complexity (at times more than 50 variants in a 50 bp window) leading to excessive memory usage (>500 GB) or disk usage (>6 TB), more details in Supplemental Information 10. To be able to handle these chromosomes, the graphs would have to be simplified prior to indexing. Indexing with CHOP yielded 103,509,254 nodes in *G^E^*, which increased the total sequence space by 14x. We again used BWA, and aligned the sample ERR052836 to both the linear reference genome and the graph, where we noted a 2-3x (13,704 to 37,826 seconds) increase in read alignment time to the graph with respect to the linear genome.

**Figure 3:**
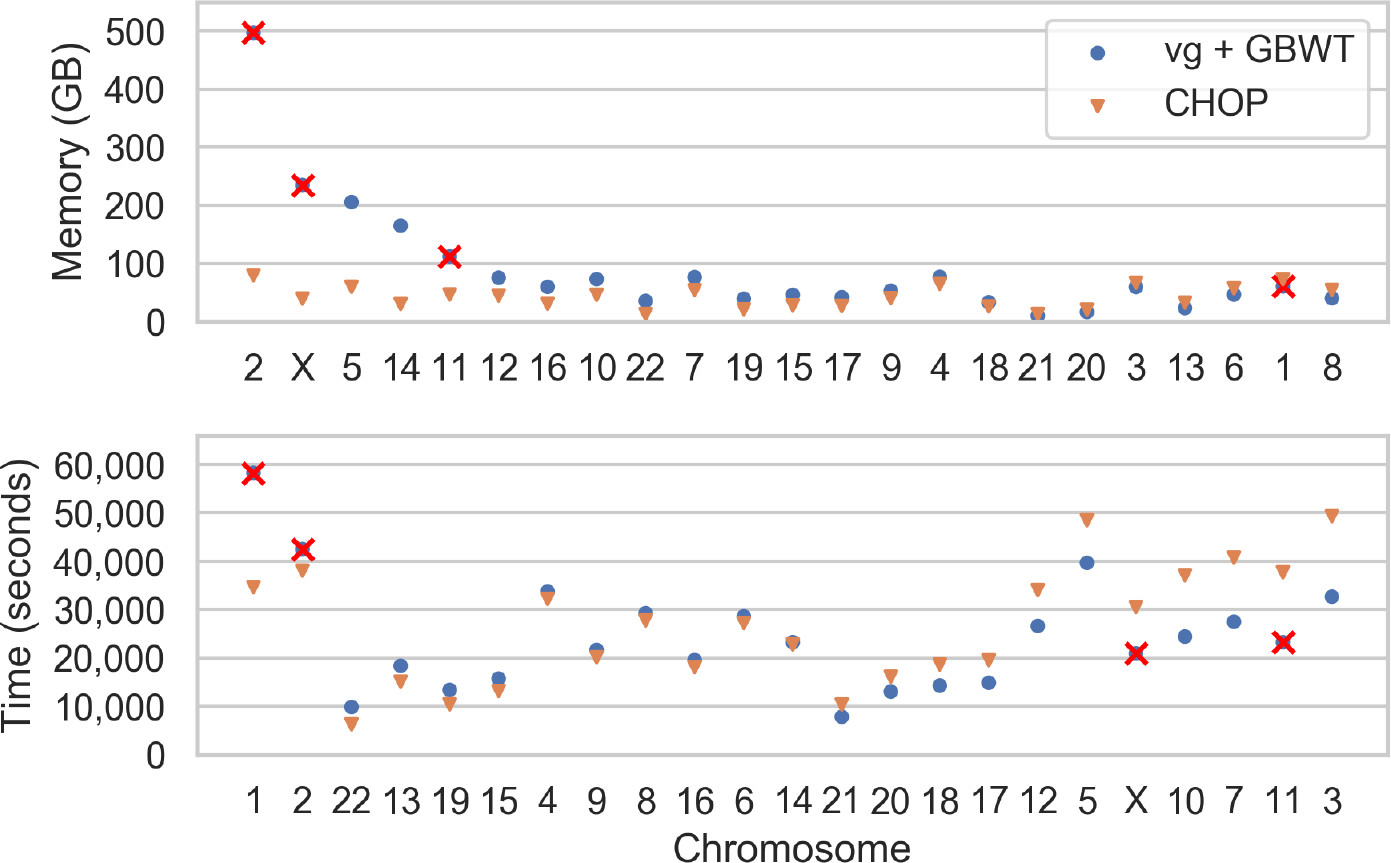
Peak memory footprint and time required for indexing the human chromosomes using CHOP and vg+GBWT. Chromosomes are ordered according to the relative differences between CHOP and vg+GBWT. Chromosomes 1, 2, 11, and X are crossed out for vg given that these ran out of memory constraints (>500 GB) or disk space constraints (>6 TB).

### 2.3 Alignment accuracy in chromosome 6

To measure alignment accuracy provided by CHOP, we compared alignments of simulated reads to multiple linear references and graph-based references. Reads were simulated using Mason 0.1.2 [29], which includes sequencing errors and base calling quality, as well as annotations denoting the ground truth location of each simulated read. Alignments are noted as correct if the aligned read is within 1 bp of the ground truth position. Only primary alignments were considered. We simulated 10,000,000 sequencing reads from a chromosome 6 sequence that encodes variants of one sample with ID NA12878. Consequently, we generated a variation set encoding only SNPs, and excluded variations and genotyping specific to NA12878 and family members.

The simulated reads were aligned to the reference chromosome 6 to provide a baseline measurement of the accuracy, as well as to a personalized chromosome 6 (linear reference including all of NA12878’s SNPs) to obtain an idealized situation. Three graphs were constructed from the NA12878 filtered variation set: Full; graph encoding all 1000G variation in chromosome 6 (excluding NA12878), Min2; graph encoding only variations that were observed in at least two individuals; PopCov10+; graph encoding the top 10% scoring variations as scored by FORGe [30], which weighs variants by allele frequency in the population and minimizes graph complexity. Figure 4a shows the fractions of reads that are correctly and incorrectly aligned onto the different reference genomes. In Figure 4b the sensitivity metrics of perfectly mapped reads and number of mismatches are shown for the same alignments.

**Figure 4:**
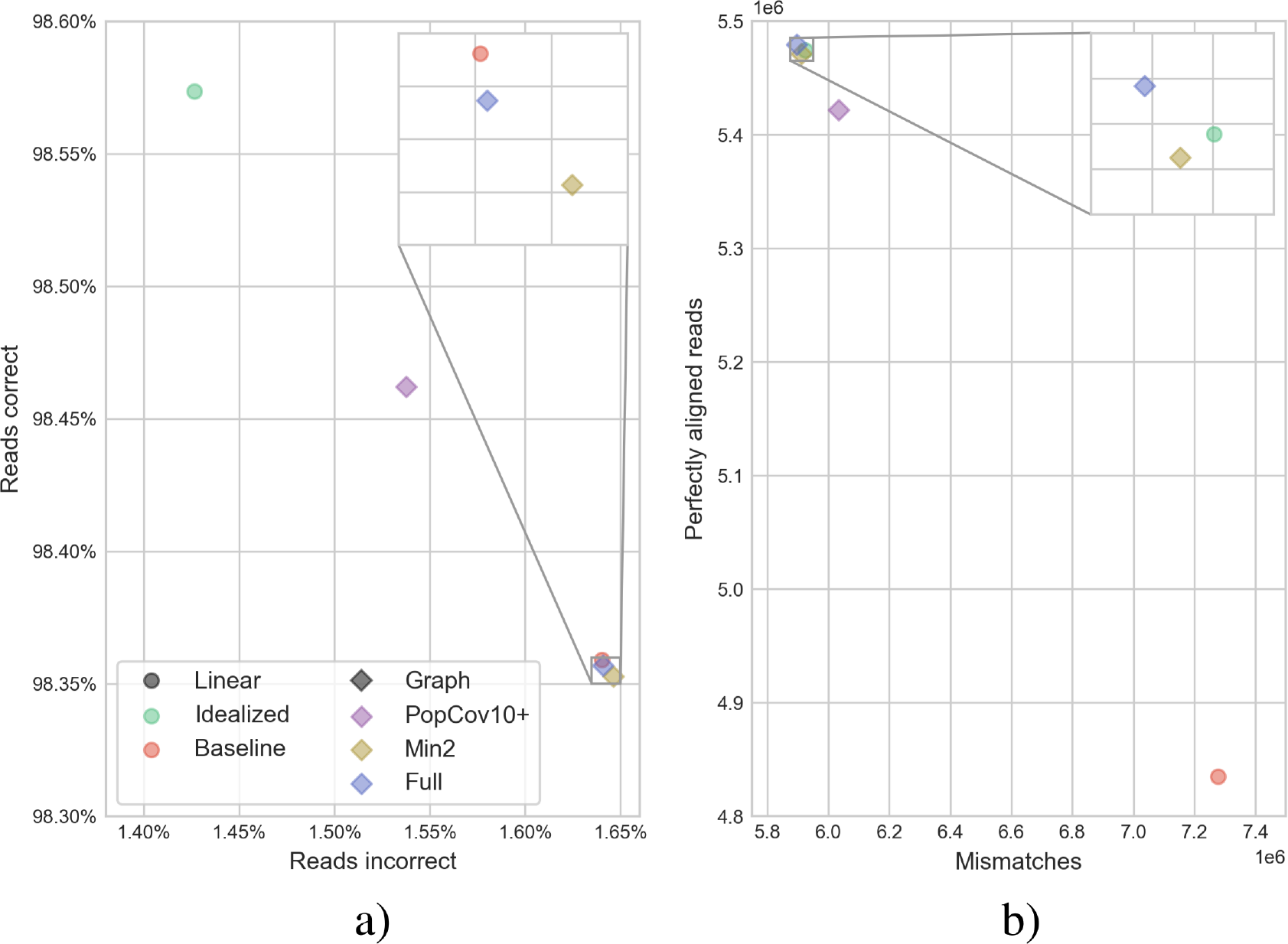
Alignment statistics of the NA12878 simulation. a) The fraction of correctly aligned reads and incorrectly aligned reads. b) Sensitivity metrics of perfectly mapped reads and number of mismatches.

Although alignment sensitivity rises as more variants are introduced into the population graphs, it also increases sequence repetitivity in the graph, which negatively influences alignment accuracy. This can be observed for both the Min2 and Full graphs which are less accurate than the baseline, while they have comparable sensitivity with respect to the idealized reference genome. The trade-off of sensitivity and specificity is clearly seen when variant selection is performed, as is the case with the PopCov10+ graph, which improves accuracy, at the cost of sensitivity.

## 3 Variation integration

As graphs span a larger search space, we investigated how this affects read alignment and variant calling. Theoretically, encoding more distinct sequences in a graph should enable alignment of more reads, and potentially allow new variants to be called. To evaluate this, variants were integrated using a feedback loop. First SRR833154 reads were aligned to H37Rv using BWA, then variants were called using Pilon. Variants were quality filtered down to 838 SNPs and then used to construct a graph with H37Rv (now thus including two genomes). The same set of reads was then aligned onto the graph, and variants were called. We expected that the additional context offered by the graph would point to previously undiscovered variants. An example of this is schematically shown in Figure 5a.

**Figure 5:**
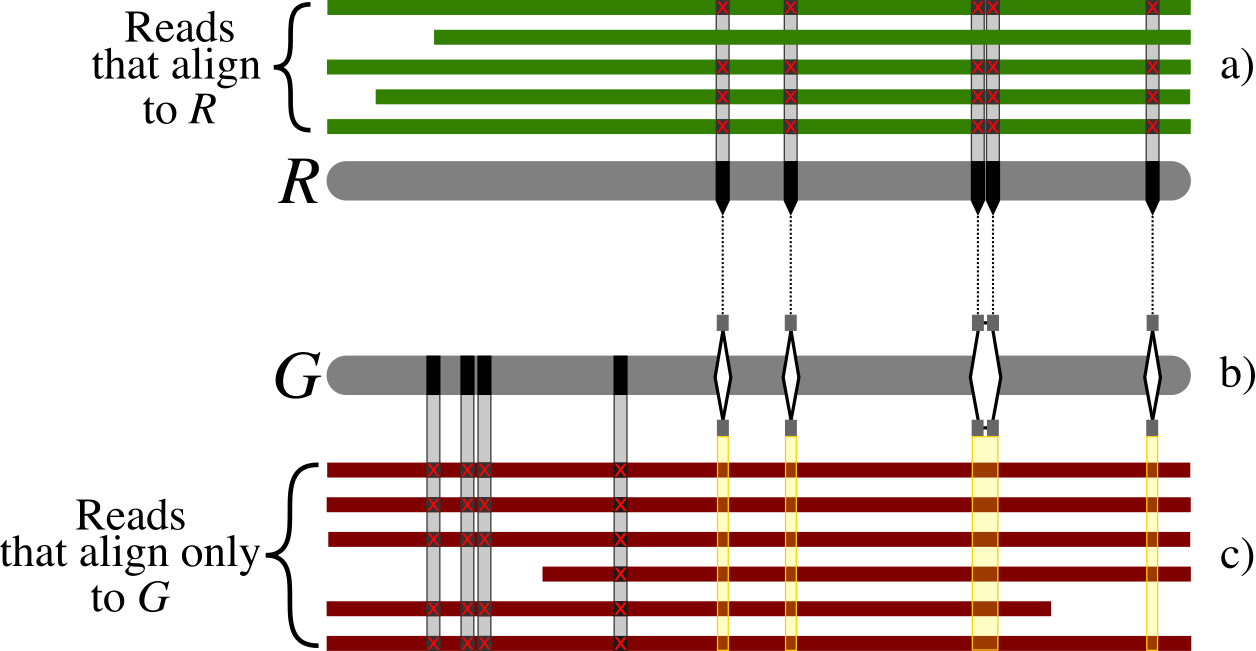
a) Schematic alignment of SRR833154 reads to the reference *R*, H37Rv, with subsequent variant calling detecting 5 high quality SNPs in this particular region. b) These and all other SNPs across the genome are integrated with the reference into graph *G*, followed by alignment of the same reads. c) Reads that previously did not align to *R*, now align onto a haplotype of the graph *G*. Formation of a pileup allows for the detection of 4 new variants in this region.

Integrating variants in a graph (Figure 5b) and realigning reads to the graph allowed reads to follow a path within the graph that best matches. This in turn allowed for reads that previously were not able to align now to be aligned. (Figure 5c). As a result, 19 (+2, 26%) new high quality variants could be called from these new aligned reads.

## 4 Discussion

Population reference graphs have the potential to improve sequencing analyses by taking into account within-species genetic diversity during the process of mapping sequencing reads. This can potentially improve various downstream analyses, like variant calling.

A challenge for mapping reads to a graph efficiently, is to find exact matching seeds of a fixed length *k* that can span the edges of the graph. Searching through an enumeration of all possible *k*-length paths in the graph is computationally challenging as the exponential growth of paths adversely affects the memory footprint as well as the alignment time. This puts practical limits on the variation that can be encoded in the population graph. We suggest the use of haplotype information to constrain this exponential growth. Doing so, the genetic linkage between neighboring variants can be exploited to counter, not only the computational problems, but also the number of false positive matches that arise due to unobserved combinations of variants (variants encoded on different alleles).

Here we introduced CHOP, a method which converts a haplotype-annotated population graph into a set of sequences that covers all observed *k*-paths. It does this by transforming the population graph into a null graph (a graph devoid of edges) such that every observed *k*-path length path is represented in one of the resulting unconnected nodes. As the resulting set of sequences (the null graph) is a compressed representation of all *k*-paths through the graph, it becomes feasible to use values for *k* that are equal to the length of typical NGS reads (e.g. 100 to 150). For this reason, an additional advantage of CHOP is that any NGS read aligner (e.g. BWA or Bowtie) can be used to map reads onto the created null graph. As every position in the null graph can be translated back to a position in the initial population graph, we can effectively perform scalable read to graph alignment.

We showed that read alignment using CHOP in combination with BWA (CHOP/BWA) easily scales to the whole human genome, encompassing all 84,745,123 variations reported by the 1000 Genomes project (2,504 individuals). The memory footprint of CHOP per human chromosome is less than 80 GB and takes under 50,000 seconds.

Furthermore, we showed that graph indexing and alignment with CHOP/BWA resulted in more aligned bases compared to aligning to the linear reference genome. Also, we found that the number of aligned bases grows proportionally with the number of incorporated variants (samples). Interestingly, the amount of sequence required to store the resulting compressed *k*-paths grew faster than the time needed to perform the alignments. We attribute this to an increase in the number of exact matching reads, which decreases the need for extending initial seeds during the alignment, which is a computational demanding task.

We extensively compared CHOP/BWA to vg, which is the current state-of-the-art toolkit for working with population graphs and includes a read alignment module. Vg uses GCSA2 indexing, an extension of the BWT for population graphs, supporting exact query lengths of up to 256 bp. Recently, vg has been expanded to facilitate haplotype constrained alignment using the GBWT, a graph extension of the positional Burrows-Wheeler transform. When comparing read alignments of CHOP/BWA with vg and vg with haplotyping (vg+GBWT) on population graphs of both *Mycobacterium tuberculosis* (MTB) and human, we found very similar alignment results, as expected. However, compared to CHOP/BWA alignment times were 5-6-times longer with vg and vg+GBWT. For analysis of human data with vg+GBWT, we had to reduce the max length of exact matching strings to 52 bp instead of the 100 bp as used in CHOP/BWA. Under those constraints, vg and vg+GBWT still exceeded the limits of our compute facility (>500GB RAM, >6TB temporary disk storage) for chromosomes with dense local variation.

Interestingly, the read alignment results did not differ much between a haplotype-constrained aligner and a non-haplotype constrained aligner. This can be best observed when comparing vg with vg+GBWT, because they utilize the same aligner and parameters, except for the haplotype pruning of all *k*-paths. Although, the number of aligned reads increases by 1.5% when considering all *k*-paths (vg) in the human population graph with respect to the linear reference genome, as opposed to an increase of 1.2% when considering haplotype-constrained *k*-paths (vg+GBWT). Inspection of the additionally mapped reads seems to indicate that most of these alignments are the result of spurious matches induced by unsupported sequence combinations. Together, this seems to suggest that considering indexing all possible *k*-paths does not add much value, while at same time increasing the chance of false positive alignments.

The advantage of limiting *k*-paths to observed haplotypes is further supported by our observation that population graph alignment improves with respect to a linear reference genome when not all observed variation is incorporated into the graph. Our simulations on a 1000G sample showed that improved read alignments (as identified by a reduced number of false positive/negative alignments) can be achieved when the allele frequency of a variant is considered when building the population graph. Simply put, if the frequency of a variant increases, it is more beneficial for read alignment to incorporate such variants in the population graph, at a minimal cost of introducing false positives. Note that rare variants within a sample can still be called after the reads are mapped to the graph, they are just not used when building the population graph.

Graphs that serve as input to CHOP should encode phased variant calls. While this information is not typically encoded in variant call formats, it is required at only short ranges (related to the value for *k*) and should be readily be available from typical sequencing experiments. In our experiments, the complexity of incorporated variation was limited to SNPs and small indels. Therefore, the benefit of a population graph on increasing the number of aligned reads was limited, since the identification of SNPs and small indels are well identifiable using a linear reference genome. However, CHOP is not restricted to graphs constructed from variant calls, but can handle any acyclic sequence graph, e.g. as generated from multi whole-genome alignments or haplotype-aware de-novo assembly algorithms [31, 32]. Consequently, both short (SNPs/indels) and long range (structural variants) haplotypes can be incorporated in the graph and in the resulting index. We expect that incorporating larger structural variations, could lead to more substantial improvements. One should realize, however, that incorporating structural variation increases the amount of repeated sequence in the graph, e.g. the incorporation of mobile element insertions and repeat expansions, which will lead to an increase in ambiguously aligned reads.

CHOP does not directly support long reads or paired reads. For long reads, with *k* typically exceeding >10 Kbp, will still lead to an intractable number of haplotype-constrained *k*-paths. However, the alignment of long reads generally depends on the detection of short seeds in the first place, which can easily be extracted from the compressed representation of *k*-paths generated by CHOP. Hence, long read alignments may be seeded, where a subgraph can be extracted (based on the seeds) and aligned to with partial order alignment. Alternatively, a heaviest weighted sub-path can be extracted from the graph [33], followed by a typical sparse-alignment on that linear sequence. For paired reads, reads are aligned to discrete *k*-paths, where an aligner such as BWA cannot directly measure distance between any distinct *k*-path. Note that read pairing should be possible based on the haplotyped paths in the graph. Namely, that the distance between any two nodes in the graph will follow a distribution of distances (of each reachable haplotype), and that this enables the evaluation of read pairs during read alignment (in a stand-alone aligner) or as a post-processing step.

Finally, we showed that by iteratively integrating mapped sequencing reads derived from one genome to the linear reference genome using the graph representation improves variant calling. Aligning additional reads led to the additional calling of variants, which subsequently could be merged with the built population graph, reiterating the whole process (multiple times). This application of population graphs is similar to iterative remapping methods, such as, ReviSeq [34], but is solved in a more general way when using population graphs as the starting point.

## 5 Methods

### 5.1 Population graph definition

Population graphs were constructed given existing reference genomes and sets of variations, called from linear reference alignments (Supplementary Information 11). Nodes within graphs are labeled, encoding genomic sequences that may be shared within multiple haplotypes, which are in turn connected by directed edges. Traversing a sequence of edges, i.e. a path, will describe either a mixture of haplotypes, or an observed haplotype within the graph.

### 5.2 Population graph specification

A population graph *G* = (*V, E*) is defined as a set of nodes *V* = *{v*_0_*, …, v_N_}*, where *N* = *|V |*, and a set of edges *E*. Each of these edges is an ordered pair of nodes (*u, v*) ∈ *E*, where node *u* ∈ *V* is connected to node *v* ∈ *V*. As *G* is a directed graph, it holds that for any edge (*u, v*) ∈ *E*, (*u, v*) ≠ (*v, u*).

For each node *v* ∈ *V*, the in-degree, *in* (*v*), is defined as the number of incoming edges to that node; i.e. the number of distinct edges (*u, v*) ∈ *E* for any *u* ∈ *V*. Conversely the out-degree of node, *v*, *out* (*v*), is defined as the number of outgoing edges from that node.

Every node, *v*, is assigned a sequence of characters, *S*, consisting of the alphabet Σ = *{A, T, C, G}*, such that *v*_*S*_ = *S* [0*, n −* 1], wherein *S* [*i*] *∈* Σ for all *i*, and the length of the sequence, *n*, is defined as *n* = *|v_S_ |*. The range of any such sequence for any node, *v* ∈ *V*, lies between 1 *≤ |v_S_ | ≤ L*, where *L* is the length of the largest recorded sequence. Any substring of a sequence, *S*, is denoted as *S* [*i, j*]. Two types of substrings in particular are prefixes *S*[0*, j*] and suffixes *S*[*i, n −* 1], which describe the left and right flanks of any sequence *S*, respectively.

A path, *P*, where *P* = *u*_0_ *…u_q−_*_1_, is any consecutive series of nodes, (*u_i_, u_i_*_+1_) ∈ *E* for all *i < q*, where *q* = *|P |* is the total number of nodes on the path. If a path exists between any pair of nodes in a graph, it is a connected graph, i.e. there are no unreachable nodes. The sequence, *S*, of a path, *P*_*S*_, is the concatenation of sequences contained in the nodes, such that *P*_*S*_ = *u*_0*S*_ · · · *u*_(*q−*1)*S*_.

Given haplotyping information, the graph *G* is augmented with a set of haplotypes, *H*, where *H* = *{H*_0_, …, *H*_*h*−1_}, where *h* = *|H|* is the number of observed haplotypes. Every edge (*u, v*) is assigned a subset of *H* denoted as (*u, v*)_*H*_, which describes the haplotypes that pass through the edge. Each encoded haplotype is represented by a path traversal through *G*, and may overlap other haplotypes.

Let *G^E^* denote the null graph of *G* such that *G^E^* = (*V*′, ∅), where *V*′ originates from merging nodes in *V* (details of which are to follow in the subsequent section).

### 5.3 Constructing the null graph

The purpose of indexing a population graph is to allow for efficient substring queries on the paths that span across nodes and edges of the graph (Figure 6). Given any non-trivial sized graph, enumerating all possible paths is often unfeasible, given the exponential nature of traversing all combinations of nodes and edges.

**Figure 6:**
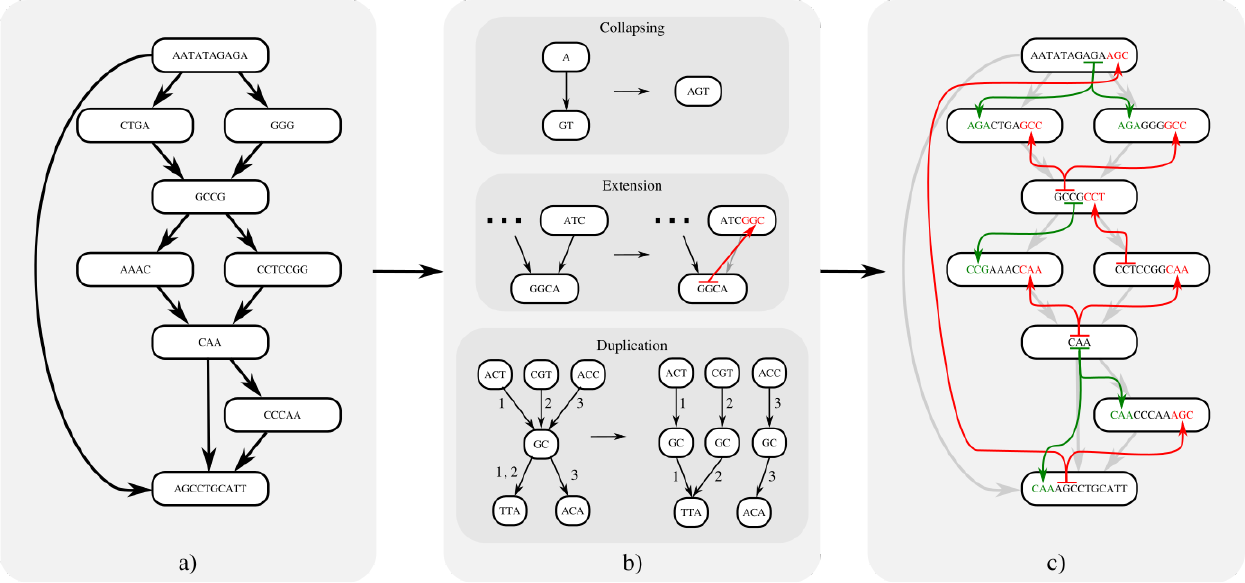
Reporting the haplotyped *k*-paths in the population graph *G* transforms it into the null graph *G^E^*, here *k* = 4. a) A population graph with sequence encodings on the nodes. b) Indexing of *k*-paths based on three operations; Collapsing, merging adjacent nodes. Extension, assigning *k*-length substrings as prefixes or suffixes between adjacent nodes. Duplication, copying of nodes and redistribution of edges among copies. c) The null graph encodes all 4-length paths in the original graph, coloring of lines and text denote the origin of assigned prefixes (green) and suffixes (red) (note that colored lines are not edges in the graph).

CHOP constrains queries through a graph to be part of a haplotype with which the population graph was built. Hereto, CHOP transforms graph *G* into a null graph *G^E^* such that every node in *G^E^* represents a sequence of length *k* or longer, and that every substring of length *k* originating from the encoded haplotypes in *G* is also a substring in a node of *G^E^*. Meaning that if sequencing reads are true error-free samplings of an underlying haplotype and are of the same length (or shorter) than the chosen value of *k*, they should correspond to a substring of a node in *G^E^*. This, in turn, enables the application of any existing read aligner to place reads onto *G^E^*. Through this transformation of *G* to *G^E^*, all haplotyped paths of at least length *k* in the graph are accounted for. The transformation is driven by three operators: collapse, extend, and duplicate (pseudocode is given in Supplemental Listing 1), explained throughout the rest of this section. While the output of CHOP can depend on the order of these three operations, we observed no significant difference in runtime or indexing outcome for different orderings.

#### Collapse

The first operation to transform *G* to *G^E^* is collapse, which merges redundant traversals of nodes in the graph. If an edge (*u, v*) ∈ *E* conforms to *out*(*u*) = 1 and *in*(*v*) = 1, then any path that traverses *u*, will immediately be followed by *v*. Therefore, it can be considered a redundant traversal such that the sequence on *u* and *v* can be merged without affecting the number of sequences that can be spelled by the graph. To do so, sequence and the corresponding intervals of *u* and *v* are merged after which the outgoing edges of *v* are transfered to *u*, followed by the removal of *v* and the edge (*u, v*). We denote this operation as collapsing, defined as *u||v* for any edge (*u, v*) (as shown in Figure 6b, pseudocode is given in Supplemental Listing 2). The direction of collapsing is guided by minimizing the number of edge reassignments, such that when *in* (*u*) *> out* (*v*), *v* is collapsed into *u*, joining the sequence *u*_*S*_ = *u_S_ · · · v_S_*. Alternatively *u* is collapsed into *v*, joining the sequence *v*_*S*_ = *u_S_ · · · v_S_*.

#### Extend

After collapsing redundant edges in the graph, a number of the remaining edges can be addressed with the extend operation. Extend is based on the observation that all *k*-length substrings that span a single edge (*u, v*), i.e. substrings that are defined by substrings of both the sequences of nodes *u* and *v*, can be accounted for by joining a *k −* 1-length substring from one node and assigning it to the other. This extension of substrings may happen bi-directionally, namely the *k −* 1-length right-hand flank of *u* is extended as a prefix of *v*, denoted as *u* ↠ *v*, provided that *in* (*v*) = 1 and *|u_S_ | ≥ k −* 1. Or vice versa, extending the *k −*1 length left-hand flank of *v* as a suffix of *u*, denoted as *u* ↞ *v*, provided that *out* (*u*) = 1 and *|v_S_ | ≥ k −* 1 (pseudocode is given in Supplemental Listing 3). To illustrate this operation consider the subgraph in Figure 7. Within this graph both nodes *u* and *v* encode sufficient sequence to allow for extension between the two and report a *k*-length overlap, resolving the edge (*u, v*). In Figure 8, a subgraph is shown in which extension is only possible for a subset of edges: (*u, w*) and (*w, v*). This does not apply for (*u, v*), as *out* (*u*) *>* 1 and *in* (*v*) *>* 1. This shows a particular situation where only after resolving nearby edges, the subgraph can be sufficiently simplified to resolve all edges. Namely, (*u, w*) and (*w, v*) must first be resolved before (*u, v*) can be solved by a collapse operation. Although the order in which substrings are extended may result in different null graphs, any of these will cover the same *k*-length substrings.

**Figure 7:**
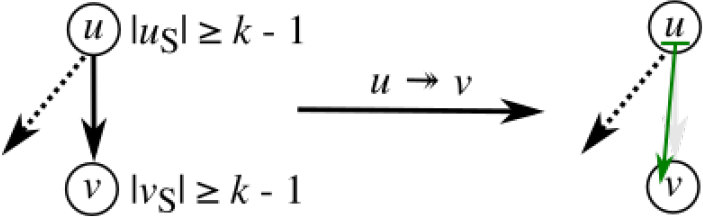
A pair of nodes *u* and *v* where |*uS*| ≥ *k* - 1 and |*vS*| ≥ *k* - 1. Note that extension is only possible by prefixing *v* with the right-hand flanking substring of *u*, given that *out* (*u*) *>* 1. The extension operation denoted as *u* ↠ *v* is defined as *v*_*S*_ = *u*_*S*_ [*|u_S_ | − k −* 1*, |u_S_ |*] *· · · v_S_*.

**Figure 8:**
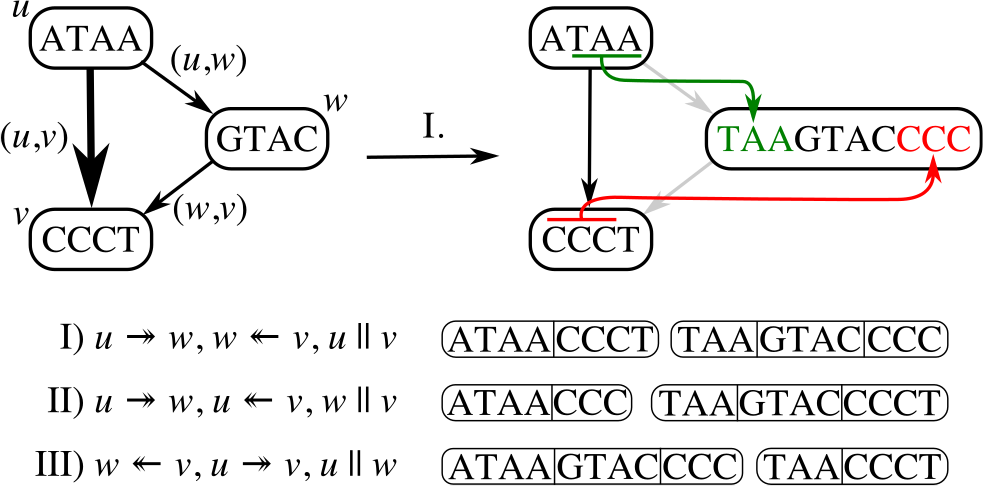
Subgraph in which substring extension for *k* = 4 between (*u, v*) is not allowed unless either (*u, w*) or (*w, v*) are resolved first. Three different solutions can resolve this subgraph, and each solution is equivalent in *k*-path space.

Since extension always concerns a *k −* 1 length prefix or suffix, any substring of length *k* that is sampled from the underlying haplotypes will exclusively correspond to either the sequence in *u* or the prefixed sequence in *v* (or vice versa). In other words, by extending and subsequently removing edges in *G* we introduce overlapping sequence as if we were converting *G* to the repeat-resolved string graph representation of a joint assembly of all genomes in *G* from all possible reads of length *k* [35].

#### Duplicate

At times neither collapse nor extend can be applied to any of the remaining edges in the graph without introducing path ambiguity, a situation in which there are multiple possible candidates to collapse or extend to/from, and choosing any candidate will block off paths to the remaining candidates. In these situations, graph topology must be simplified through the third operation, duplicate. The duplicate operation duplicates a node such that the set of incoming and outgoing edges are split between the duplicated nodes. (pseudocode is given in Supplemental Listing 4). Duplication allows consequent collapsing, which in turn enables substring extension, such that after a sufficient number of iterations all edges in *G* can be resolved, either by means of extension or collapsing.

As opposed to methods that aim to track all possible paths through the graph, we suggest the use of haplotype information that is modeled on the edges to constrain the number of necessary node duplications from *in* (*u*) *∗ out* (*u*) to *δ*. Where *δ* is the number of paired incoming and outgoing edges for *u* that have at least one intersecting haplotype. Note that *δ* is bounded by the number of haplotypes encoded in *G*, and that there will never be more duplications than haplotypes in any one region of the graph.

To illustrate the idea, Figure 9 shows a subgraph with haplotypes encoded on the edges. From the haplotyping we can derive that not all paths through this graph are supported by the underlying haplotypes. For example, the path *u → d → f* combines sequence segments that are unsupported (the haplotypes between (*u, d*) and (*d, f*) do not overlap). By excluding these unsupported paths through the graph, the number of duplications for node *d* can be constrained from 6 to 3. This way, the search space for subsequent *k*-length substrings is greatly reduced with respect to reporting all possible paths. Supplemental Figure S1 gives the full details about the transformation from Figure 1a to Figure 1b.

**Figure 9:**
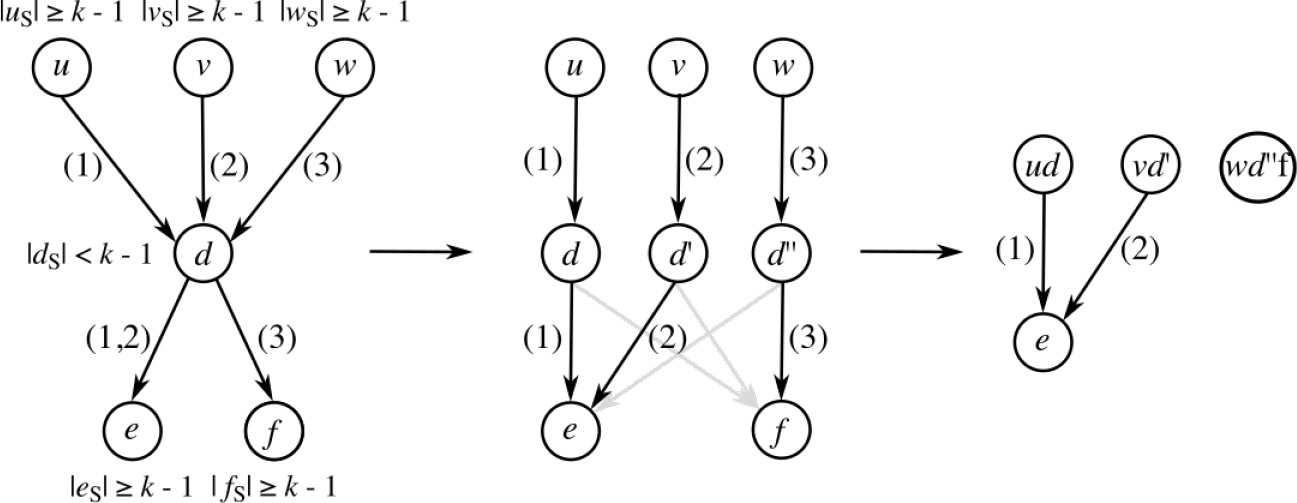
Subgraph with haplotypes: {1, 2, 3}. Node *d* must be duplicated, as no more edges can be removed through extension or collapsing without introducing ambiguity. By grouping incoming and outgoing haplotypes on *d*, the number of duplications can be reduced. In the resulting graph, edges (*u, d*), (*v, d′*), and (*w, d′′*) can be collapsed. Finally, an extension can be applied to edges (*ud, e*) and (*vd′, e*) which would lead to the null graph. Note that the introduction of grayed out edges is prevented using haplotyping, hence the edge count is reduced from 6 to 3.

### 5.4 Mapping reads to CHOP’s null graph

Established alignment tools can now be used to directly align reads to the null graph representation as long as reads are shorter or equal to *k* + 1. Because the sequence modeled on the nodes in *G^E^* is now a composition of sequence originating from adjacent nodes in *G*, the intervals that gave rise to these compositions need to be traced in order to convert the alignment of a read to a node in *G^E^* to a path in *G*. For this reason, during the transformation from *G* to *G^E^*, the originating node in *G* and corresponding offset for each prefixed, suffixed or concatenated sequence is stored alongside the actual sequence. Note that in theory the defined operations can also be expressed purely in terms of interval operations, excluding any sequence. Given the intervals, a mapping between *G^E^* and *G* is maintained, such that any node in *G^E^* can be traced back to the corresponding path of nodes in *G*. As a result, any alignment to a node in *G^E^* can also be traced to a sub-path of this path, effectively enabling the alignment of reads to graph *G* by using *G^E^* as a proxy (Figure 1c).

