## Supplemental information for "CHOP: Haplotype-aware path indexing in population graphs"

### Supplemental materials

#### 1 Transformation to a null graph

CHOP can through consecutive steps of extension, collapsing, and duplication (as described in the methods) transform population graphs into null graphs. In this edgeless graph representation each node now describes a $k$-length path through the original graph. In Supplemental Figure 1, we describe how the graph in Figure 1a is transformed into the null graph of Figure 2b.

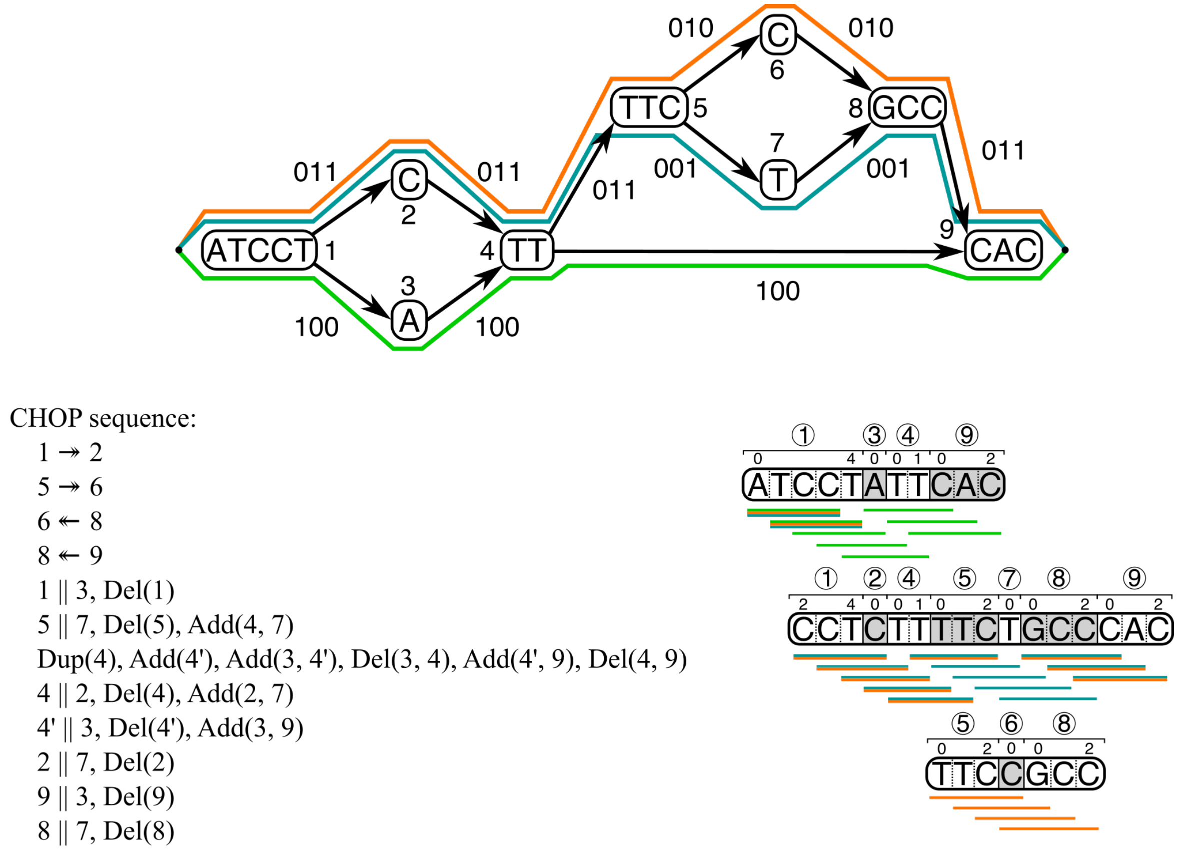


Figure 1: The same graph as in Figure 1a, now shown with haplotypes on the edges encoded as bitvectors. Using CHOP, the input graph can be transformed into a null graph. Each of the steps performed by CHOP are shown in sequential order; Extension: $x \twoheadrightarrow y$ ($y$ is prefixed by $x$), and $x \twoheadleftarrow y$ ($x$ is suffixed by $y$). Collapsing: $x||y$ ($x$ and $y$ are collapsed into a single node). Duplication: Dup($x$), (node $x$ is duplicated).

#### 2 *Mycobacterium tuberculosis* read sets

We obtained the 10 holdout samples from EBI-ENA, as shown in Table 1. All reads have a length of 101 bp, and are single-end.

Table 1: Samples used in TB experiments, associated read sets are included, with KRITH1/2 accession numbers.

| Sample | Read set | KRITH1/2 ID | Read count |
|---|---|---|---|
| TKK-01-0053 | SRR833154 | G28639 | 5,263,942 |
| TKK-04-0029 | SRR1019154 | G47382 | 5,481,779 |
| TKK-02-0022 | SRR1011463 | G47310 | 4,301,550 |
| TKK-01-0093 | SRR958234 | G38246 | 9,249,605 |
| TKK-02-0066 | SRR924236 | G32253 | 5,168,899 |
| TKK-01-0016 | SRR832997 | G27617 | 8,615,425 |
| TKK-02-0051 | SRR847783 | G32041 | 7,571,230 |
| TKK-01-0039 | SRR833147 | G27616 | 6,443,482 |
| TKK-01-0047 | SRR832984 | G27644 | 5,532,779 |
| TKK-01-0033 | SRR833024 | G27582 | 7,582,870 |

### 3 Variation growth in *Mycobacterium tuberculosis* population graphs

When constructing graphs for the hold-out experiment, progressively more samples (from 1 to 400) are included in the constructed graphs. By including more samples, more variants are incorporated into the graphs, as can be seen in Figure 2.

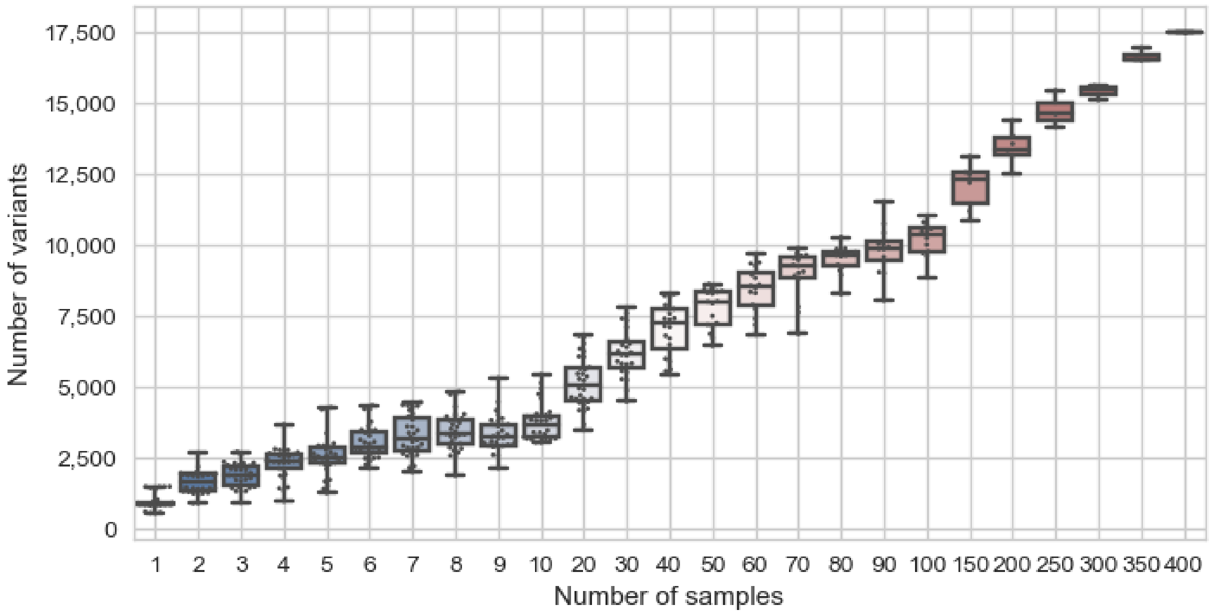


Figure 2: Variable size variant sampling for VCF-based graph construction.

### 4 Alignment criteria used for evaluation

To evaluate behavior of different aligners we measure the following criteria: number of mismatches, insertions, deletions, clipped bases, aligned reads/bases, unaligned reads/bases, perfectly aligned reads, and non-primary alignments. Mismatches, insertions, deletions, and base clipping may all be introduced to allow the alignment of reads onto the reference. To handle substitutions between reference and query, mismatches are introduced. Multiple base pair divergences are treated as insertions to the reference (query) or as deletions from the reference (reference). Base clipping masks portions of reads (from either end) that do not align to the reference from end to end, meaning shorter but contiguous read fragments are aligned. The extent of these operations in alignment can particularly characterize differences in alignments to linear references and population graphs. With the expectation that the incidence of these operations decreases in graph alignments (in proportion to the number of aligned bases).

The number of aligned and unaligned reads are indicative of the proportion that aligns in a read set. For instance reads may not be aligned at all because of insufficient sequence context on the reference or due to low read quality, random noise, and/or contamination. The number of bases that are aligned provide more detail, given that not all reads are perfectly aligned. Perfectly aligned reads, describe full length alignments of reads for which no mismatches/insertions/deletions/clipping are introduced. The number of bases that are unaligned includes the bases from unaligned reads, mismatches, insertions, and clipped bases. Reads for which there are multiple valid alignments that score equally, result in non-primary alignments. Meaning that for every read there will always be a primary alignment (or it is unaligned), and one or more non-primary alignments. The incidence of these non-primary alignments give an indication on the extent of ambiguity in the alignments, given that this is typically induced by repetitivity in the reference.

### 5 Comparing VCF-based population graphs of CHOP and vg

Because the graph construction methods of CHOP and vg are similar but not the same and this can also affect indexing (see Figure 3), we tested whether the read alignment to the two different graphs differed. For that we aligned reads from sample SRR833154 with the vg aligner to graphs from the hold-out experiment using either vg or CHOP. We measured the perfectly aligned reads, unaligned reads, and mismatches where we took the ratio between these measures when aligning against the graph constructed by vg and CHOP. A ratio of 1 implies no difference. The results in Figure 4 show that there is minimal difference.

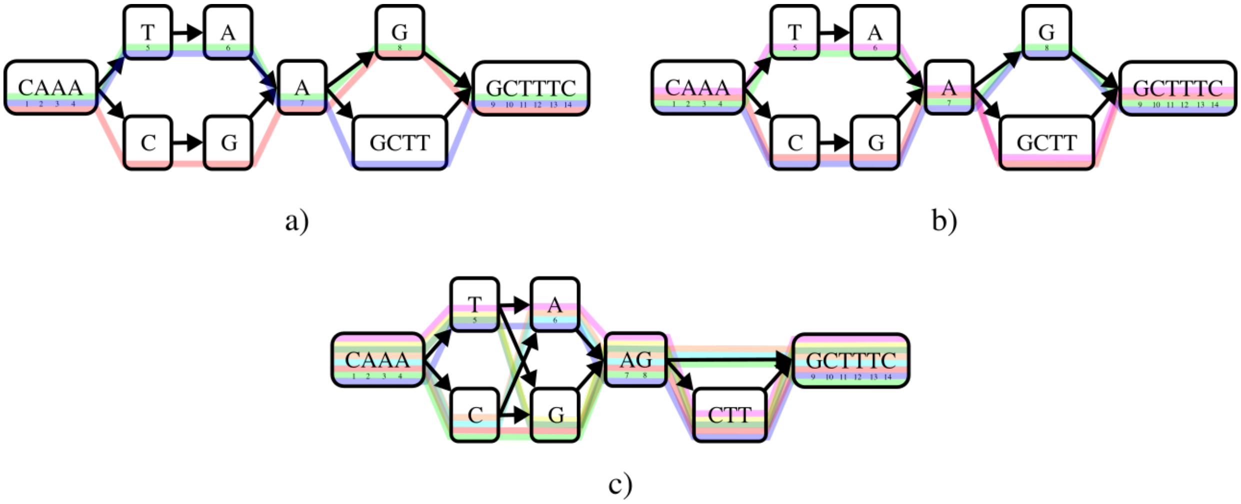


Figure 3: Graph construction of CHOP and vg (Supplemental Figure 12) can affect the resultant paths that are indexed. a) CHOP extracts two paths from the CHOP constructed graph. b) If the same graph is indexed with vg, there are four paths. c) The vg constructed graph indexed by vg has eight paths.

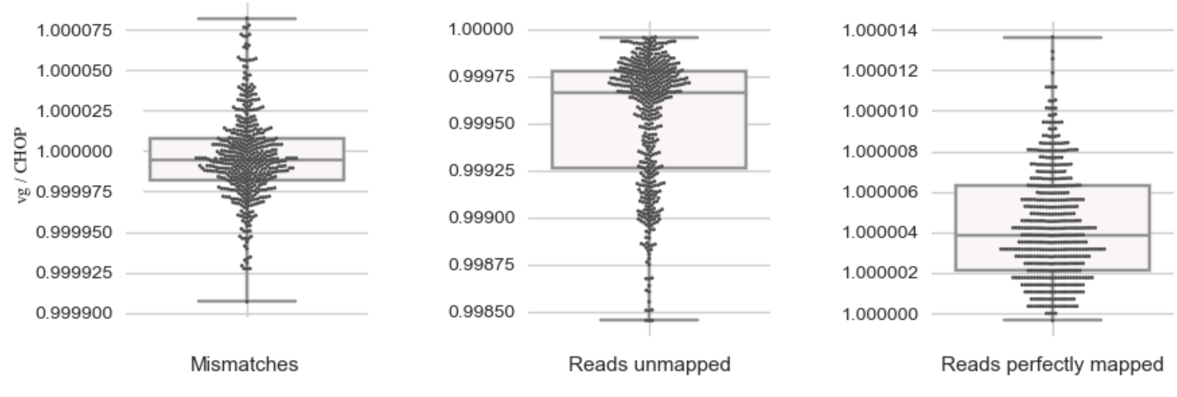


Figure 4: SRR833154 alignments with vg using graphs constructed from both CHOP and vg. The ratio (vg over CHOP) for mismatches, unaligned reads, and reads that are perfectly aligned.

### 6 *Mycobacterium tuberculosis* read alignment to H37Rv and graph

In the hold-out experiment 10 different single-end read sets are aligned to the reference genome, H37Rv, and to population graphs that progressively include more samples (up to 400 excluding the hold-out). Alignments were evaluated on both a read and base-count basis. Shown for SRR833154 this includes the number of perfectly aligned reads (Figure 2), unaligned reads (Figure 5), and mismatched bases (Figure 6).

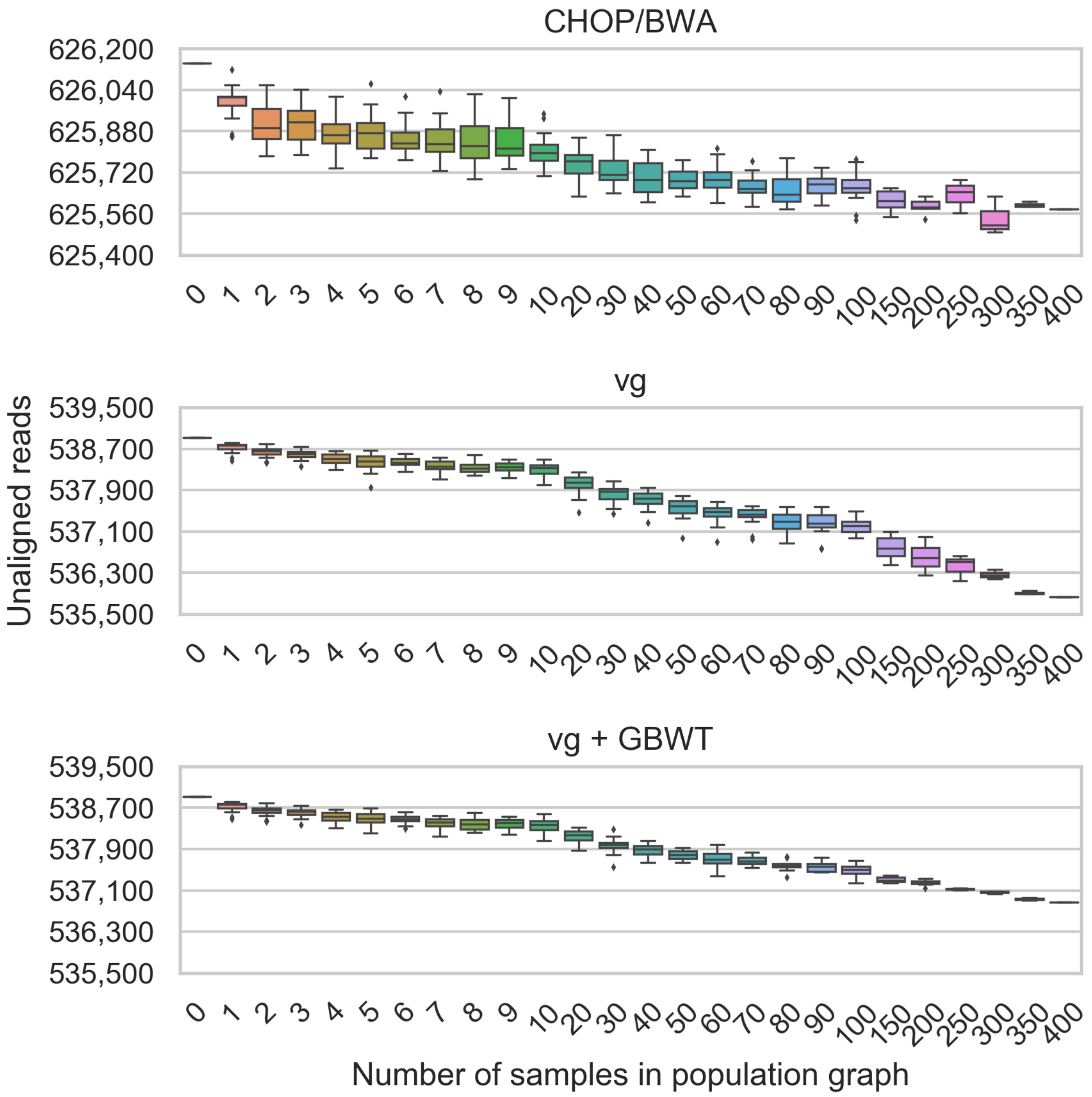


Figure 5: Unaligned read count for SRR833154 alignments to different sized population graphs, containing between 0 (only H37Rv the linear reference) and 400 samples.

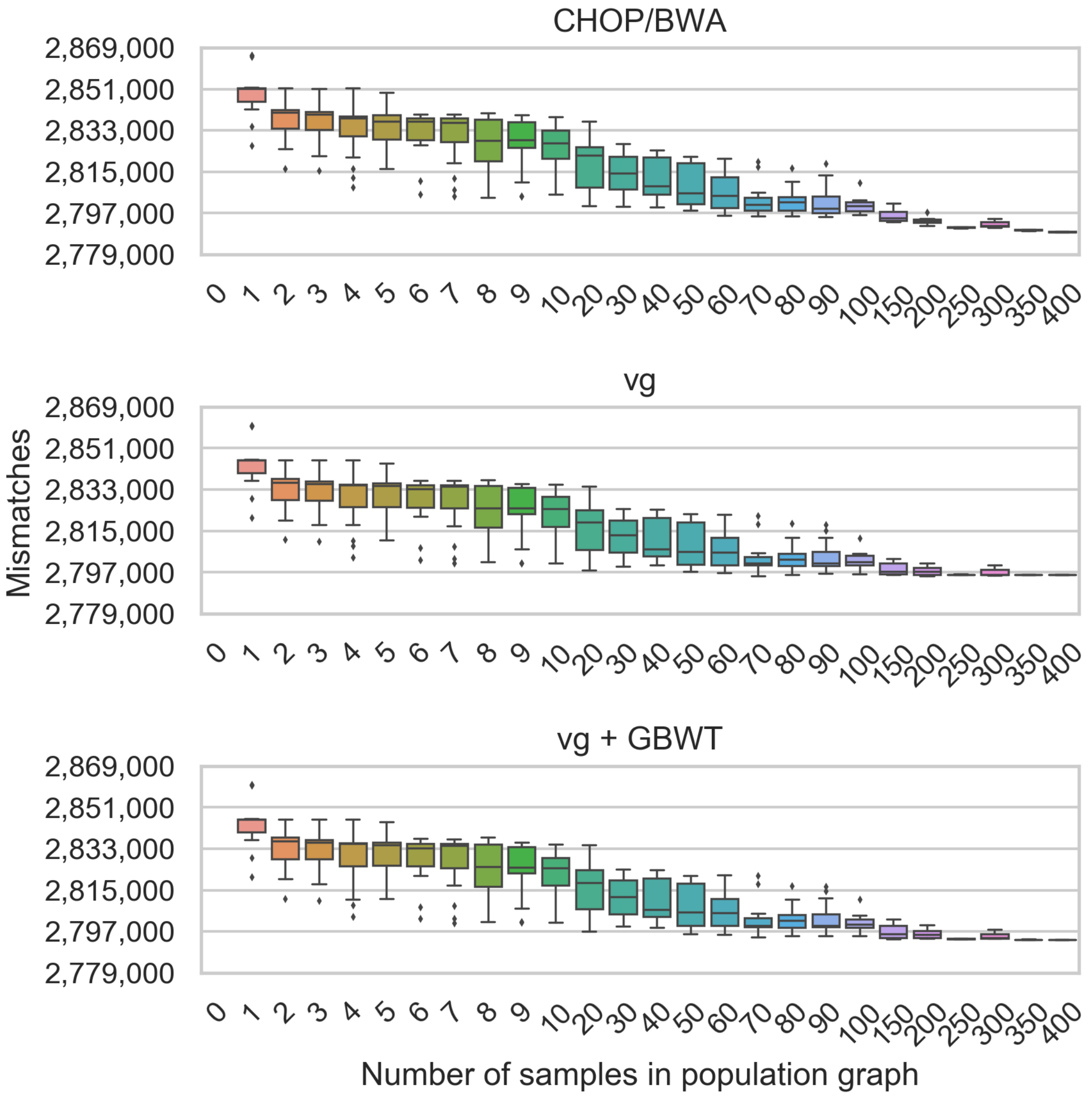


Figure 6: Mismatch base count for SRR833154 alignments to different sized population graphs, containing between 0 (only H37Rv the linear reference) and 400 samples.

### 7 Variation linkage of 1000 Genomes in chromosome 6

The 1000 Genomes Phase 3 variant set of chromosome 6 encodes 5,023,970 variants. In order to determine how much of the variation was shared among samples, we evaluated the genotyping of every variant as is shown in Figure 7. This revealed that 41.58% of all variants are unique to its sample.

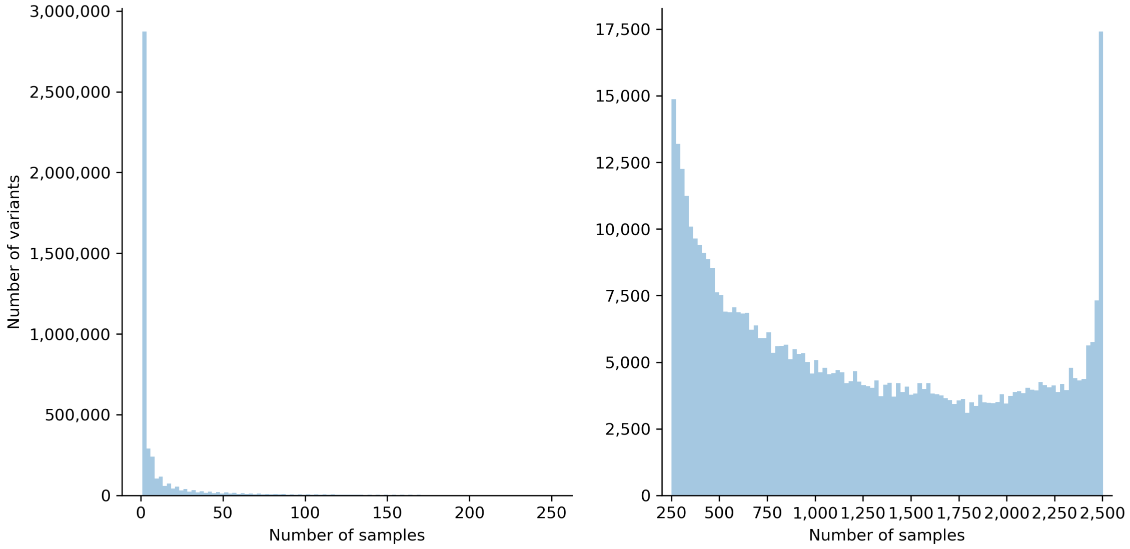


Figure 7: Two histograms displaying the extent of shared variations among samples in the 1000 Genomes Phase 3 data.

#### 8 Filtered 1000 Genomes read sets

Since we align only to a graph of chromosome 6, the single-end read sets (Table 2) were first filtered to exclude any reads aligning to other chromosomes. This was accomplished by aligning all 15 read sets to the human genome (excluding mitochondrial DNA) using BWA, and subsequently generating new read sets by extracting reads that were aligned to either chromosome 6 or those that were unaligned.

Table 2: The read sets from the 1000 Genomes used in alignments to chromosome 6.

| | | | Filtered reads | | |
|---|---|---|---|---|---|
| Population | Sample | Read set | Count | Mapped to Chr6 | Unmapped |
| ESN | HG02938 | ERR257960 | 6,238,375 | 5,572,661 (89.33%) | 665,714 (10.67%) |
| ESN | HG03521 | ERR257962 | 6,012,874 | 5,252,933 (87.36%) | 759,941 (12.64%) |
| FIN | HG00308 | ERR050084 | 3,882,577 | 2,814,118 (72.48%) | 1,068,459 (27.52%) |
| FIN | HG00380 | ERR050085 | 4,234,048 | 3,280,314 (77.47%) | 953,734 (22.53%) |
| GBR | HG01791 | ERR052834 | 3,066,482 | 2,358,096 (76.90%) | 708,386 (23.10%) |
| GBR | HG01789 | ERR052836 | 3,454,819 | 2,664,211 (77.12%) | 790,608 (22.88%) |
| GIH | NA20881 | ERR068420 | 2,278,409 | 2,015,089 (88.44%) | 263,320 (11.56%) |
| GIH | NA20884 | ERR068423 | 1,765,696 | 1,545,396 (87.52%) | 220,300 (12.48%) |
| IBS | HG01670 | ERR050090 | 1,115,839 | 859,298 (77.01%) | 256,541 (22.99%) |
| IBS | HG02223 | ERR056986 | 1,808,334 | 1,467,485 (81.15%) | 340,849 (18.85%) |
| KHV | HG01595 | ERR059932 | 1,217,726 | 1,059,586 (87.01%) | 158,140 (12.99%) |
| KHV | HG02017 | ERR059937 | 1,375,752 | 1,205,911 (87.65%) | 169,841 (12.35%) |
| MSL | HG03054 | ERR251326 | 4,720,003 | 4,293,293 (90.96%) | 426,710 (9.04%) |
| MSL | HG03378 | ERR251401 | 3,650,563 | 3,263,964 (89.41%) | 386,599 (10.59%) |
| YRI | NA18517 | ERR239432 | 569,541 | 478,520 (84.02%) | 91,021 (15.98%) |

#### 9 Reads mapping to mitochondrial DNA

Figure 8 displays the distribution of the number of aligned bases for reads that aligned exclusively onto the graph.

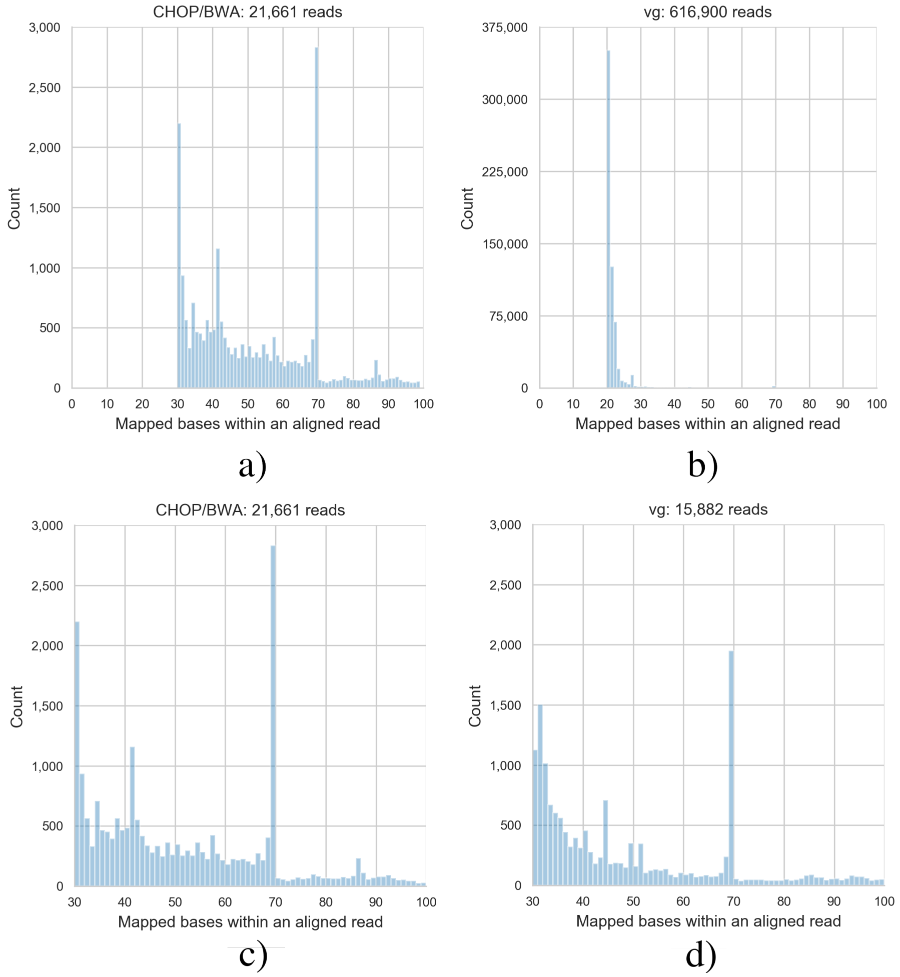


Figure 8: The number of reads that have a particular number of bases aligned after their alignment onto the chromosome 6 population graph with CHOP/BWA (a) and vg (b), respectively. In c) and d) the same is shown in the range of 30 to 100 bases.

Of the reads corresponding to the peak at 69 bp as shown in Figure 8, 97.54% of them aligned to the same fragment of a path in the graph. We used BLAST ( [1]) to determine the origin of the path and found hits on chromosome 6 and mitochondrial DNA (corresponding to the fragment of the path). Realigning the same reads to mitochondrial DNA revealed that most of the reads map full length (100M), as is partially shown in the pileup of Figure 9.

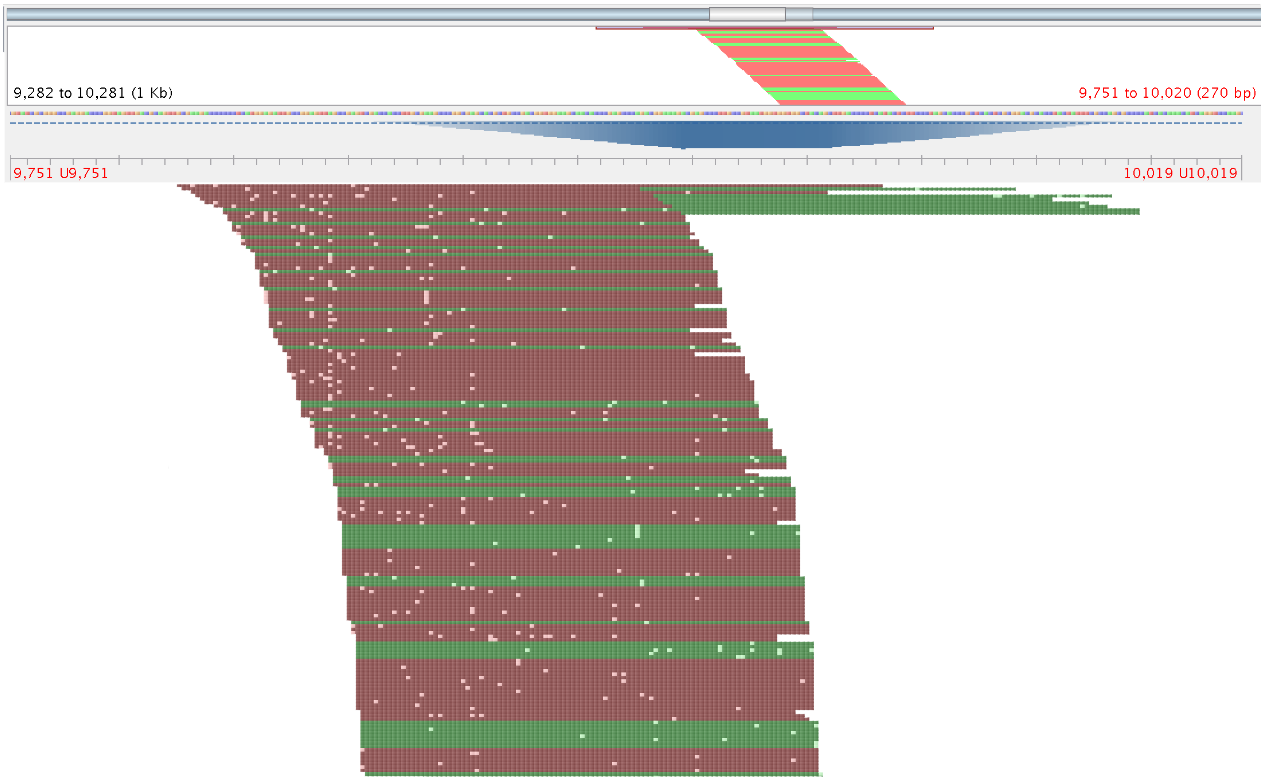


Figure 9: Pileup visualization of mitochondrial DNA using Tablet of reads that were previously unmappable on the linear reference genome (excluding mitochondrial DNA) but mappable on the graph that actually correspond to mitochondrial DNA ( [2]).

### 10 Variant density in human chromosomes

The computational costs required to index human chromosomes is highly dependent on the number of variants encoded in the graph as well as the density of these variants and the chromosome size. For certain human chromosomes this variation density (possibly in combination with chromosome size) can lead to an explosive growth in the required memory/disk space, which is what happened with vg+GBWT for chromosomes 1, 2, 11, and X. To illustrate this, we quantified the number of variants across each chromosome in windows of 50 bp (note that we set vg to index $k = 52$ length paths) as is shown in Figure 10. The chromosomes 1, 2, 11, and X each encode variants at higher densities than others, and chromosome 1 even exceeds 50 variants in a 50 bp window, note that these measurements should also be put into context of the chromosome size.

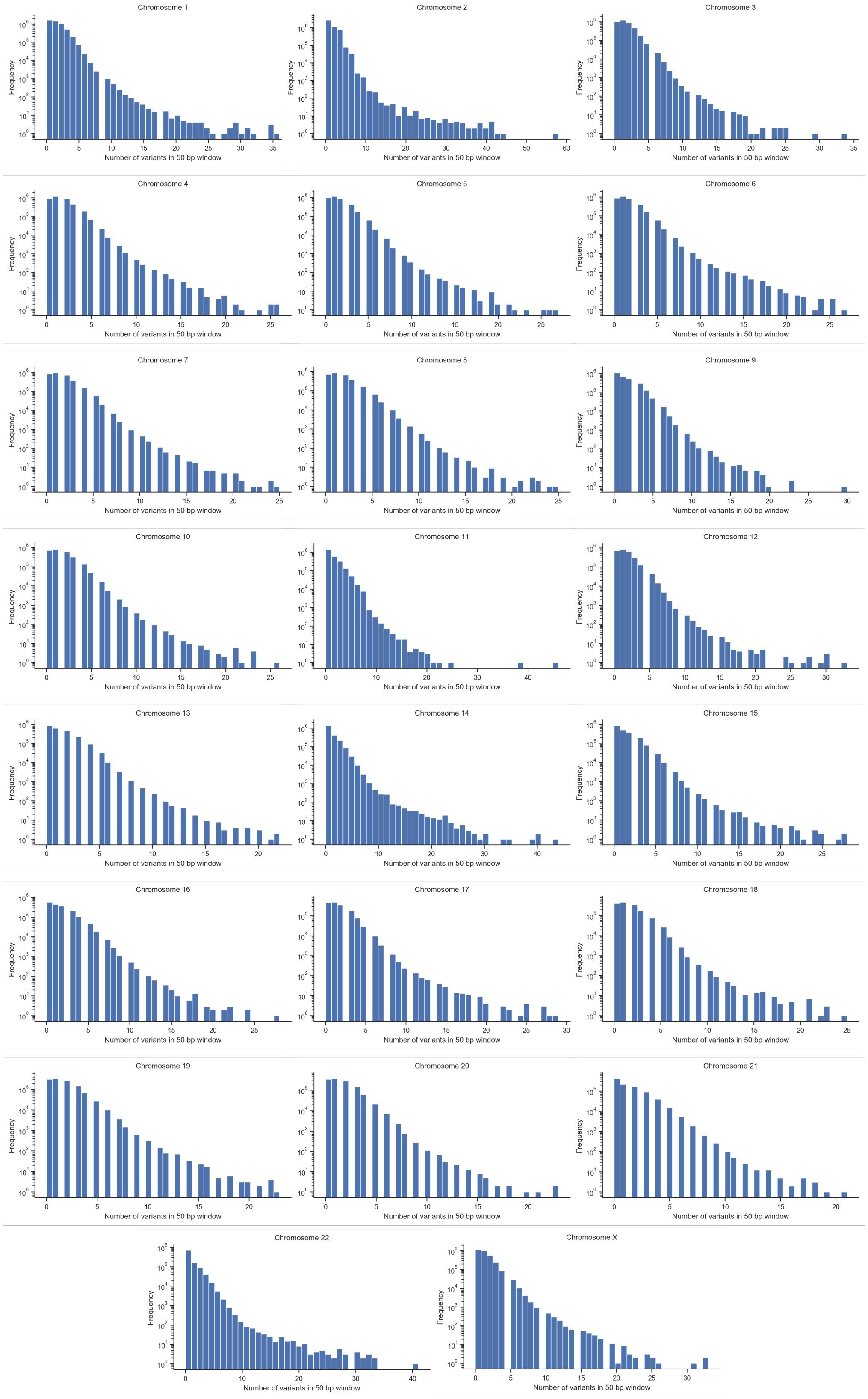


Figure 10: Variant distribution in 50 bp windows for each human chromosome.

### 11 Constructing a population graph from known variants

One way of constructing population graphs, is projecting sets of variations (from VCF files) called with respect to a reference back onto this reference (Similar to construction in others methods such as vg and Graphtyper) (Figure 11a). Initially a singleton graph is created, which encodes the reference sequence (Figure 11b). Variants are, according to their reference coordinate ordering, iteratively inserted in the graph. For each variant, a minimum of three nodes is introduced into the graph. The reference node is first split into two nodes, describing sequence before and after the variation. Between these reference nodes the reference and alternate alleles are introduced (Figure 11c). In case of consecutive variants (variations at most one base-pair apart), the reference and variant alleles are connected to the preceding nodes and only then converge into a reference node (Figure 11d). The same procedure applies for both indels and SNPs when introducing variants into the graph (Figure 11e). Haplotyping information is embedded on the edges, which includes both the samples and the reference.

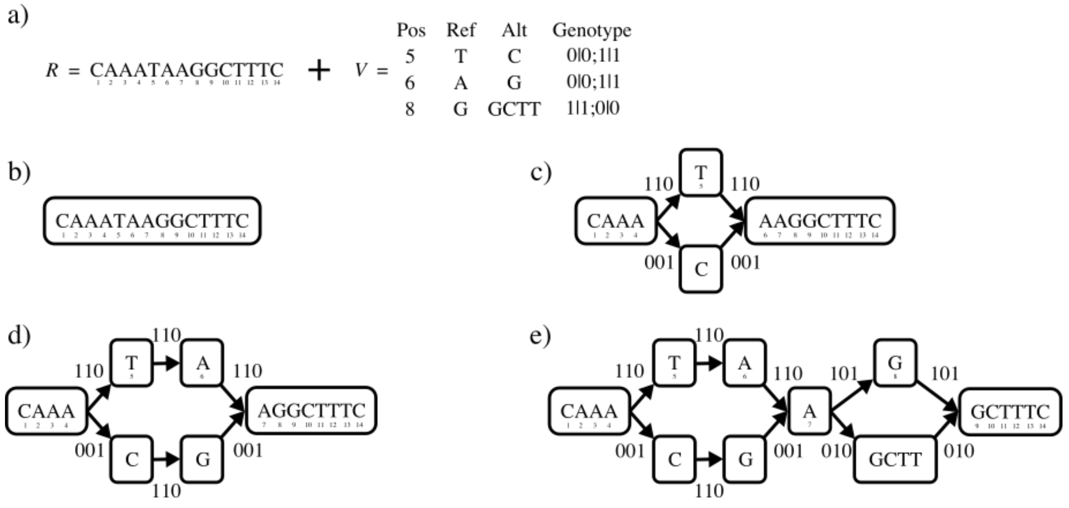


Figure 11: a) The reference sequence and variation set used in graph construction. b) A graph is initialized with a single node encoding the reference sequence. c) In order of the coordinate space of the reference, variant T → C is introduced into the graph. d) A consecutive variant is added to the graph. e) An insertion is added to the graph. The final population graph encodes three paths, one of which being the reference path.

The described graph construction strategy of CHOP differs to that of vg. The most notable change being the full combination of (consecutive) alleles and treatment of SNPs and indels, as is shown in Figure 12 for CHOP and vg.

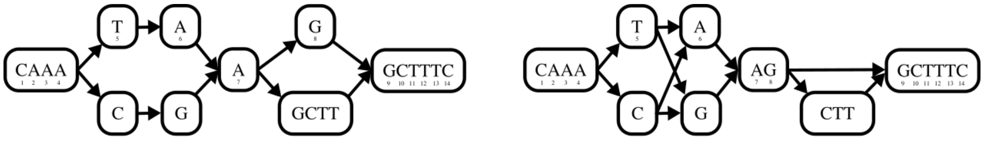


a) b)

Figure 12: Graph construction with the same input genome and variants as in Figure 11. a) Graph construction using CHOP. b) Graph construction with vg construct.

#### 12 Pseudocode CHOP procedures

**Algorithm 1**

```
procedure ChopGraph(G)
    SimplifyGraph(G)                          ▷ Extend and Collapse until exhaustion
    if G_E ≠ ∅ then
        for each edge (u, v) ∈ G_E do
            if deg(u) > deg(v) then
                Duplicate(G, u)                ▷ Duplicate u
            else
                Duplicate(G, v)                ▷ Duplicate v
        ChopGraph(G)

procedure SimplifyGraph(G)
    modified ← True
    while modified do
        modified ← False
        for each edge (u, v) ∈ G_E do
            if out(u) = 1 and in(v) = 1 then
                Collapse(G, u, v)              ▷ Collapse u||v
                modified ← True
            else if in(v) = 1 and |u_S| ≥ k − 1 then
                Extend(G, u, v, 1)             ▷ Prefix u ↠ v
                modified ← True
            else if out(u) = 1 and |v_S| ≥ k − 1 then
                Extend(G, u, v, 0)             ▷ Suffix v ↞ u
                modified ← True
```

**Algorithm 2**

```
procedure Collapse(G, u, v)
    if in(u) > out(v) then                     ▷ u ← v
        u_S = u_S · · · v_S                    ▷ Concatenate sequence
        for each edge (v, x) ∈ G_E do          ▷ Outgoing edges u
            add edge (u, x)
        delete node v
    else                                       ▷ u → v
        v_S = u_S · · · v_S                    ▷ Concatenate sequence
        for each edge (x, u) ∈ G_E do          ▷ Incoming edges v
            add edge (x, v)
        delete node u
```

**Algorithm 3**

**procedure** EXTEND$(G, u, v, isPrefix)$
  **if** $isPrefix$ **then** ▷ $u \twoheadrightarrow v$
    $v_S = u_S\left[|u_S| - k - 1, |u_S|\right] \cdots v_S$
  **else** ▷ $v \twoheadleftarrow u$
    $u_S = u_S \cdots v_S\left[0, k - 1\right]$
  **delete edge** $(u,\ v)$

**Algorithm 4**

**procedure** DUPLICATE$(G, u)$
  **for** each edge $(p, u) \in predecessors\,(G, u)$ **do**
    **for** each edge $(u, s) \in successors\,(G, u)$ **do**
      group $\leftarrow (x, u)_H \cap (u, x)_H$
      **if** group $\neq \emptyset$ **then**
        **create node** $i$ ▷ $i \leftarrow u$
        **create edges** $([(p, i)\,, (i, s)])$
  **delete node** $u$
